# CDK1 facilitates RAD51-mediated DNA repair to protect dictyate stage arrested oocytes from genotoxic stress

**DOI:** 10.64898/2026.06.27.734932

**Authors:** Ajith Kumar E, Lava Kumar. S, Pravin Birajdar, Akshay Kumar, Anjali Kumari, P. Kiran Kumar, Mohd Athar, Aradhana Mohanty, Ankita Verma, L Pradnya, G Sowmya, M Rajender, S Abhilasha, Sahina Sabnam, Partha Sarathi Nial, Y Silambresan, H.B.D. Prasada Rao

## Abstract

Oocytes arrested at the dictyate stage of meiosis I must maintain genomic integrity for prolonged periods to preserve female fertility. During this extended arrest, DNA lesions arising from endogenous and exogenous sources threaten oocyte survival, yet the molecular mechanisms coordinating DNA repair in dormant oocytes remain poorly understood. Here, we identify cyclin-dependent kinase 1 (CDK1) as a critical regulator of the oocyte DNA damage response and homologous recombination (HR) repair under genotoxic stress. Using cisplatin-induced DNA damage models in fetal goat ovaries and neonatal mouse ovaries, we investigated repair mechanisms operating within the ovarian reserve. Label-free proteomic profiling revealed significant enrichment of DNA damage response pathways following cisplatin exposure, with CDK1 emerging as one of the most prominently upregulated kinases. Pharmacological inhibition of CDK1 had little effect on follicle survival under physiological conditions but aggravated oocyte and follicle loss following DNA damage, indicating a stress-dependent role for CDK1 in preserving ovarian follicle pool integrity. Mechanistically, DNA damage activated a Chk2-dependent signaling pathway that promoted p63 phosphorylation and altered the WEE1–CDK1 regulatory axis, resulting in reduced inhibitory CDK1 phosphorylation (Thr14/Tyr15) and increased activating phosphorylation (Thr161). Activated CDK1 was associated with enhanced RAD51 phosphorylation and accumulation at DNA damage foci, supporting homologous recombination (HR)-mediated repair in dictyate-arrested oocytes. In contrast, CDK1 inhibition reduced phospho-RAD51 levels, impaired RAD51 localization, increased persistent γH2AX accumulation, and elevated oocyte apoptosis. Notably, suppression of CDK1 was accompanied by increased expression of the non-homologous end joining (NHEJ) marker Ku80 and the nucleotide excision repair (NER) factor XPA, suggesting increased engagement of alternative DNA repair pathways. Furthermore, inhibition of Chk2 abolished the DNA damage-associated CDK1 activation signature and restored WEE1 expression, supporting a model in which CDK1 functions downstream of Chk2 signaling during the oocyte DNA damage response. Collectively, our findings identify a previously unrecognized Chk2–CDK1–RAD51 signaling axis that coordinates homologous recombination repair in dormant oocytes and safeguards ovarian follicular pool integrity under genotoxic stress. These findings provide new mechanistic insight into how dictyate-arrested oocytes maintain genome stability during prolonged meiotic arrest.

## Introduction

The dictyate stage arrest represents a critical oocyte development phase characterized by prolonged dormancy during prophase I of meiosis(1). This arrest, which extends from fetal development until ovulation, plays a crucial role in preserving the genetic integrity of oocytes for decades(1). During this extended period of dormancy, the oocytes must maintain their viability and quality by effectively managing and repairing any DNA damage that may arise(2). The ability to detect and repair DNA damage is vital for ensuring oocyte health, as any deficiencies in these processes can lead to impaired fertility and potential transmission of genetic anomalies(2,3).

DNA damage is an inherent biological phenomenon that can arise from both endogenous and exogenous sources(2,4). Endogenous factors include errors during DNA replication and oxidative stress, while exogenous factors encompass environmental pollutants, chemotherapy, and radiotherapy(4,5). The DNA of ovarian cells, encompassing both oocytes and somatic cells, is particularly susceptible to damage due to their complex roles in meiosis, follicular development, and hormonal regulation. Such damage can lead to a range of reproductive and health issues, including accelerated ovarian ageing, decreased fertility, and an increased risk of ovarian diseases such as cancer(6,7). If DNA double-strand breaks (DSBs) are unresolved, apoptosis is essential to prevent germline mutations(7). The p53 family members, including p53, p63, and p73, play significant roles in regulating oocyte apoptosis in response to DNA damage, with TAp63α isoform being a key mediator(2,8–10). TAp63α, highly expressed in primordial follicle oocytes, triggers apoptosis by activating pro-apoptotic Bcl-2 family members like PUMA and NOXA(8,11). As oocytes transition from the resting primordial pool to secondary follicles, TAp63α expression decreases, potentially explaining why growing oocytes are less susceptible to DNA damage-induced apoptosis, indicating the crucial role of TAp63α, Chk2, and PUMA in DNA damage-induced apoptosis and preserving oocyte health(3,11,12). In primordial follicle oocytes, DNA damage rapidly activates the ATM/ATR signaling cascade, leading to phosphorylation of histone H2AX (γH2AX) and activation of checkpoint kinases including Chk2. Chk2 subsequently phosphorylates the oocyte-specific transcription factor TAp63α, a master regulator of oocyte quality control that determines whether damaged oocytes undergo repair or apoptotic elimination. Genetic studies in mice have demonstrated that disruption of the ATM–Chk2–TAp63α pathway can partially rescue oocyte loss following irradiation or chemotherapy, highlighting the importance of checkpoint signaling in regulating ovarian reserve integrity.

Although substantial progress has been made in understanding checkpoint-mediated elimination of severely damaged oocytes, much less is known about the mechanisms that actively repair DNA lesions in dormant oocytes before irreversible apoptosis is initiated. Recent research indicates that homologous recombination (HR) is likely the primary mechanism for repairing DSBs in primordial follicle oocytes(13,14). RAD51, a crucial HR protein, is present in these oocytes, while BRCA1 and BRCA2 are essential for effective DSB repair(15,16). BRCA1 deletion leads to diminished reproductive capacity by reducing the number of primordial follicles and increasing DSBs, whereas BRCA2 deletion results in embryonic lethality(17,18). Women with BRCA1 mutations show reduced serum AMH levels and diminished ovarian reserve, which correlate with poorer ovarian stimulation responses and earlier onset of menopause(19). Despite these insights, the precise mechanisms and critical factors involved in DNA repair within dictyate stage arrested oocytes, particularly in response to endogenous and exogenous DNA damage, remain fully understood.

CDK1 is classically recognized as the master regulator of M-phase entry and meiotic resumption in oocytes. During dictyate arrest, CDK1 activity is tightly suppressed through inhibitory phosphorylation at Thr14 and Tyr15 mediated by the kinases WEE1 and MYT1, thereby preventing premature germinal vesicle breakdown and meiotic progression(20–22). Upon hormonal stimulation, activation of the cyclin B–CDK1 complex drives chromosome condensation, nuclear envelope breakdown, and spindle assembly required for meiotic maturation(20). Beyond its canonical role in cell cycle regulation, increasing evidence from mitotic systems indicates that CDK1 also contributes to DNA repair processes, particularly homologous recombination. CDK1-dependent phosphorylation has been shown to regulate multiple HR-associated proteins, including BRCA1, CtIP, and RAD51, thereby coordinating DNA repair with cell cycle progression(23–27). However, whether CDK1 participates in DNA repair within dormant dictyate-stage oocytes remains unknown.

## Results

### Establishing DNA damage in goat ovaries to elucidate the underlying DNA repair mechanisms

To investigate the DNA repair mechanisms in dictyate-arrested oocytes within the ovarian reserve, we utilized late embryonic goat ovaries (crown-rump length: 30 cm), which provide several advantages over murine models due to their rich population of dictyate-stage oocytes (Fig. 1C and Fig S1A-C)(3,28,29). The larger size of goat ovaries facilitates the collection of greater quantities of oocytes and enhances experimental reproducibility. Additionally, goat ovaries better mimic the physiological environment of human ovaries, making them more relevant for translational research(29,30).

**Figure 1.**
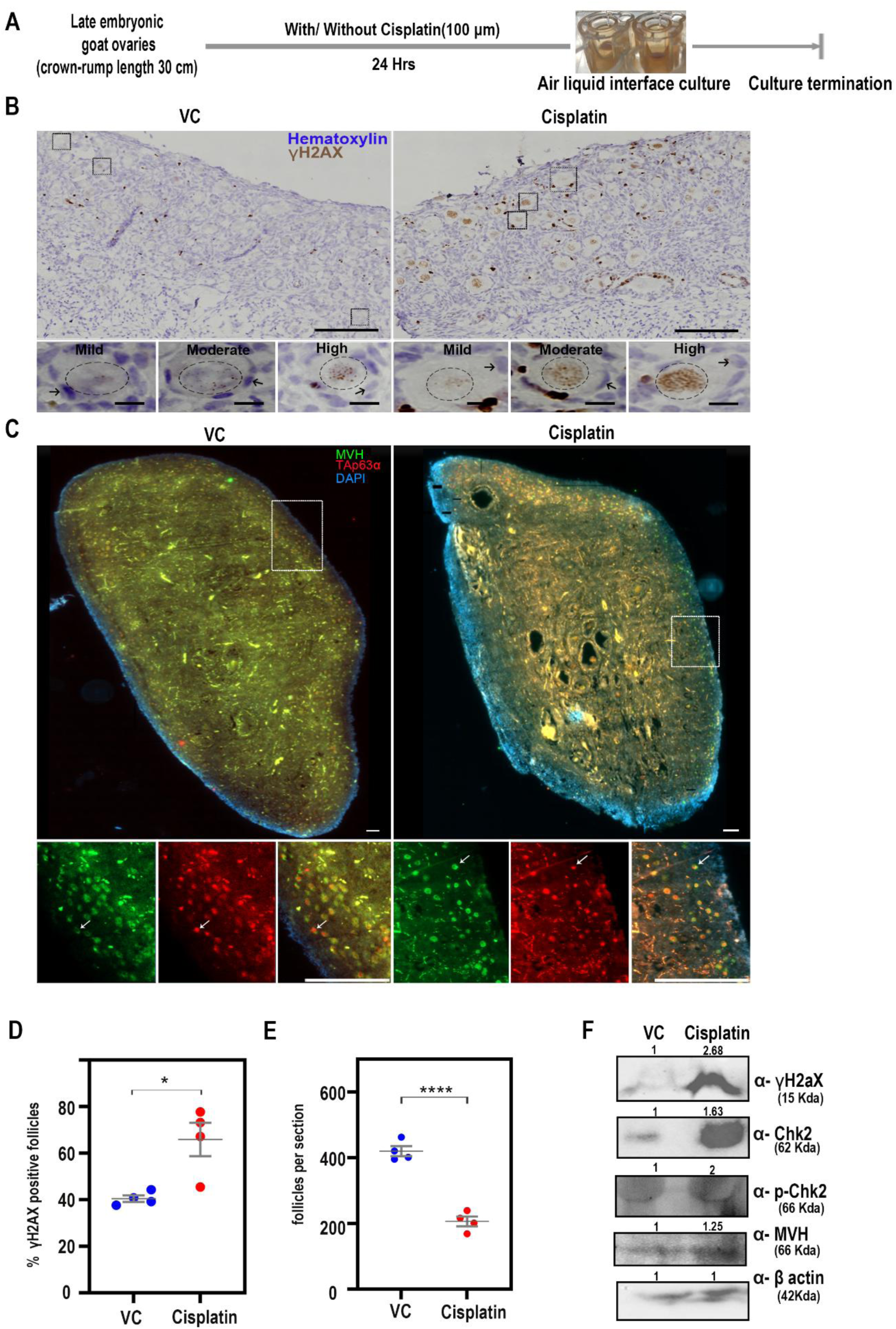
Development of a cisplatin-induced DNA damage model in cultured goat ovaries. (A) Schematic overview of the experimental approach used to induce DNA damage in late embryonic goat ovaries cultured ex vivo using cisplatin. (B) Representative immunohistochemical images of ovarian sections from vehicle control (VC) and cisplatin-treated groups stained for γH2AX, a marker of DNA double-strand breaks. Sections were counterstained with hematoxylin. Black squares indicate regions shown at higher magnification in the lower panels. Black arrows indicate granulosa cells, and black dotted circles outline oocyte nuclei. Scale bars: 100 μm (upper panels) and 10 μm (lower panels). (C) Representative immunofluorescence images of ovarian sections from vehicle control (VC) and cisplatin-treated groups stained for MVH (green), TAp63α (red), and DAPI (blue). White squares indicate regions shown at higher magnification in the lower panels. Arrows indicate follicles positive for TAp63α expression. Scale bars: 100 μm (upper panels) and 20 μm (lower panels). (D) Quantification of γH2AX-positive follicles per section in vehicle control (VC) and cisplatin-treated ovaries (n = 4 sections per group). (E) Quantification of total follicle numbers per section in vehicle control (VC) and cisplatin-treated ovaries (n = 4 sections per group). (F) Representative western blot analysis of ovarian lysates from vehicle control (VC) and cisplatin-treated groups showing expression of γH2AX, Chk2, phospho-Chk2 (p-Chk2), MVH, and β-actin as a loading control. Data are presented as mean ± SD. Statistical significance was determined using an unpaired two-tailed Student’s *t*-test. Significance levels are indicated as ****P ≤ 0.0001 and *P ≤ 0.01.

To determine the optimal concentration of cisplatin that induces substantial DNA damage while preserving a significant proportion of the ovarian reserve, goat ovaries were cultured at an air-liquid interface and treated with varying concentrations of cisplatin, a well-established DNA-damaging agent, for 24 hours (Fig. 1A)(31). Following treatment, ovarian sections were immunostained with γH2AX (a marker of DNA double-strand breaks, H2AX phosphorylated at serine 139), or MVH (a germ cell marker) and TAp63α (a key regulator of oocyte quality and survival), to quantify follicle integrity and assess DNA damage(Fig. 1B and C)(32,33).

Quantitative analysis demonstrated that treatment with 100 μM cisplatin for 24 hours resulted in a strong γH2AX signal and a 1.2-fold increase in γH2AX-positive follicles compared to the control group, while maintaining approximately 50% follicle survival (Fig. 1D and E). These results establish this treatment condition as optimal for downstream assays. Complementary Western blot analysis of protein extracts from treated ovaries confirmed a 3-fold increase in γH2AX levels relative to controls (Fig. 1F). Furthermore, increased expression of Chk2 and its activated form, phosphorylated Chk2 (pChk2), confirmed activation of the DNA damage response pathway, indicating that the selected treatment effectively models DNA damage within the ovarian reserve (Fig. 1F)(2).

### CDK1 is upregulated in early follicle oocytes in response to cisplatin-induced DNA damage

We performed label-free proteomic profiling on control and cisplatin-treated goat ovaries to elucidate the differential regulation of the DNA repair proteome during ovarian reserve DNA repair mechanisms(34). A total of 4992 proteins were identified in control ovaries and 4971 in cisplatin-treated ovaries (n = 3). Among these, 175 proteins were uniquely identified in the control group, and 154 proteins were uniquely identified in the cisplatin-treated group, while 4817 proteins were common to both samples (Fig. 2A). Notably, approximately 38% of the samples exhibited significant separation between the control and cisplatin-treated ovarian lysate in multivariate discriminative analysis (Fig. 2B).

**Figure 2:**
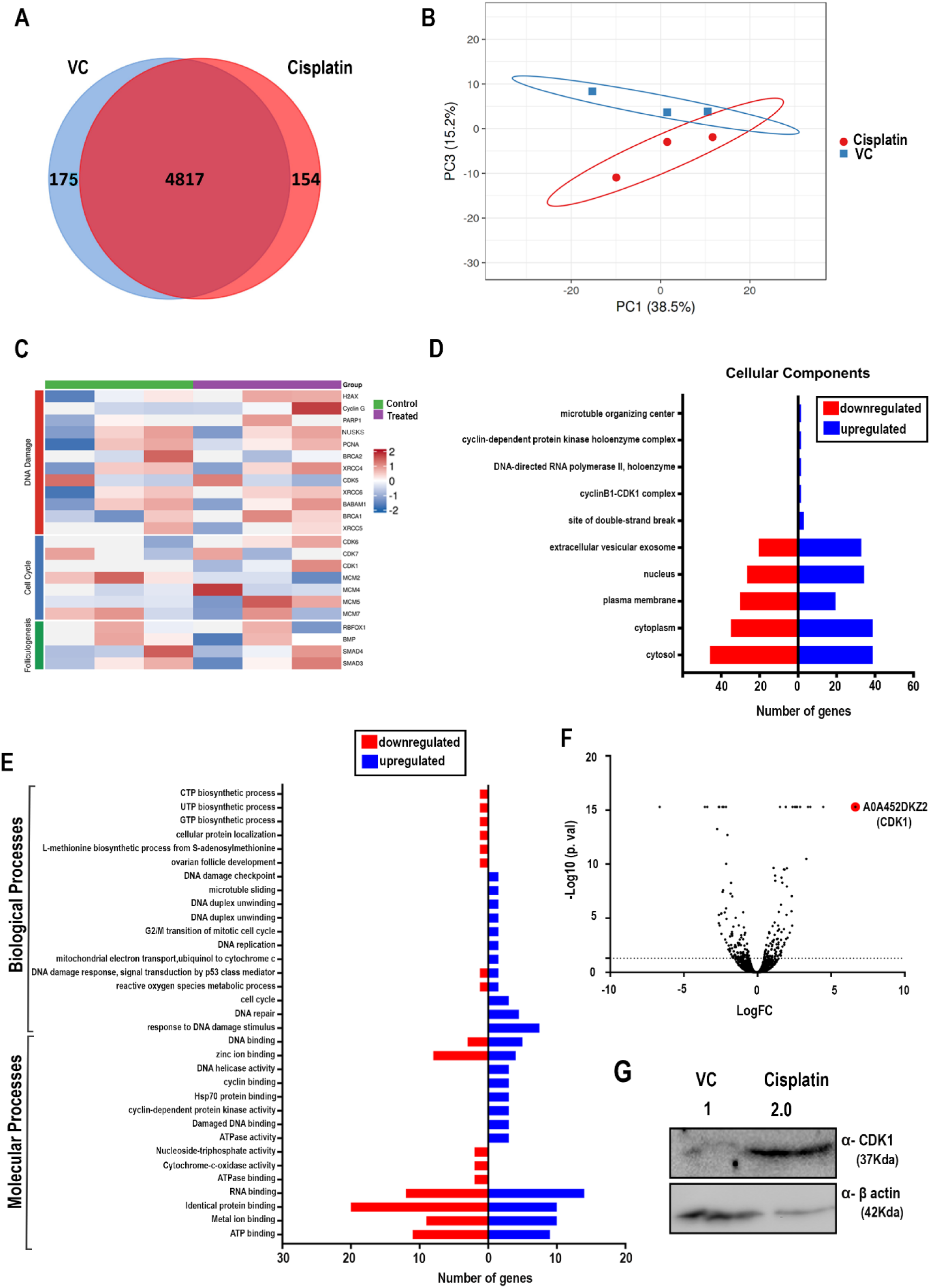
Differential proteomic analysis of vehicle control (VC) and Cisplatin-treated ovaries. (A) Venn diagram illustrating the overlap and distinct proteins identified in the proteomes of vehicle control (VC) and Cisplatin-treated ovaries. (B) Principal Component Analysis (PCA) demonstrating variance and clustering patterns between vehicle control (VC) and Cisplatin-treated ovary proteomes. (C) Heatmap depicting the differential expression of selected proteins associated with DNA damage and repair, cell cycle progression, and folliculogenesis in vehicle control (VC) and cisplatin-treated ovaries, highlighting pathway-specific changes between groups. (D, E) Gene Ontology (GO) enrichment analysis was conducted to assess overrepresented cellular components, biological processes, and molecular functions among the differentially expressed proteins. (F) Volcano plot visualizing significantly upregulated and downregulated proteins in Cisplatin-treated ovaries compared to vehicle control (VC). (G) Representative western blot images showing expression levels of CDK1 and β-Actin in VC and Cisplatin-treated ovary samples.

A heat map was generated for all identified proteins, and hierarchical clustering analysis using Euclidean distance and Ward’s linkage revealed two distinct clusters corresponding to the control and cisplatin-treated ovaries (Fig. 2C, Fig.S2). Classification of the identified proteins based on their biological functions indicated that pathways related to cell cycle progression, DNA repair, and responses to DNA damage stimuli were significantly upregulated in cisplatin-treated ovaries. Conversely, proteins associated with ovarian follicular development and reactive oxygen species (ROS) metabolic regulation were downregulated compared to controls (Fig. 2D and E).

Molecular functional pathway analysis further indicated an upregulation of cyclin-dependent protein kinase (CDK) activity in cisplatin-treated ovaries, warranting a detailed examination of protein abundance differences (Fig. 2E). Among the top 50 most abundant proteins, we observed a notable upregulation of CDK1 in cisplatin-treated ovaries (Fig. 2F). This finding was validated by Western blot analysis, which confirmed a 2-fold increase in CDK1 expression following cisplatin-induced DNA damage (Fig. 2G).

### CDK1 activity is dispensable for maintaining the ovarian follicle reserve under physiological conditions

To evaluate whether CDK1 is essential for maintaining the natural ovarian follicle reserve, we conducted both in vitro and in vivo experiments using goat ovarian cultures and neonatal mice, respectively.

### In Vitro Study (Goat ovarian cultures)

Goat ovarian tissues were cultured and treated with the CDK1 inhibitor RO-3306 at concentrations of 250 μM and 500 μM, or with PBS as a vehicle control, for up to 24 hours (Fig. 3A)(35). RO-3306 is a known ATP-competitive inhibitor that blocks the kinase activity of CDK1. After treatment, the ovarian tissues were fixed, sectioned, and immunostained for MVH (a germ cell marker) and TAp63α (a regulator of oocyte integrity) (Fig. 3B). Follicle counts were performed on the stained tissue sections (Fig. 3C). The results showed no significant difference in follicle numbers between the RO-3306-treated and control groups, suggesting that CDK1 inhibition may not impact follicle maintenance under these in vitro conditions (Fig. 3C).

**Figure 3.**
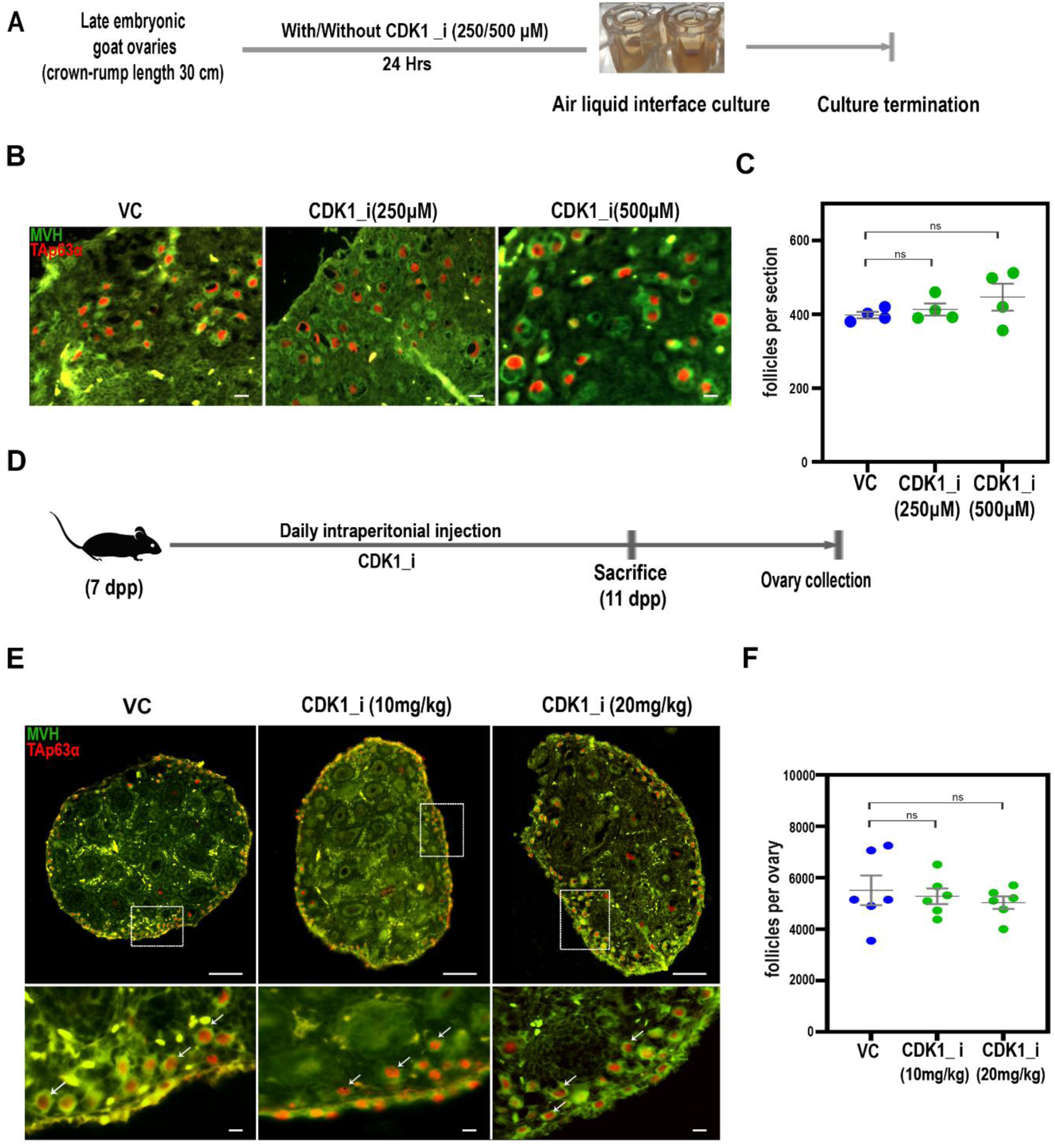
Pharmacological inhibition of CDK1 does not affect maintenance of the ovarian follicle pool. (A) Schematic overview of the experimental design used to assess the effects of CDK1 inhibition in cultured late embryonic goat ovaries. (B) Representative immunofluorescence images of ovarian sections from vehicle control (VC) and CDK1 inhibitor-treated (CDK1i) goat ovaries stained for MVH (green) and TAp63α (red). Scale bar = 50 μm. (C) Quantification of total follicle numbers per ovarian section in vehicle control (VC) and CDK1 inhibitor-treated goat ovaries (n = 4 sections per group). (D) Schematic overview of the in vivo experimental design showing intraperitoneal administration of a CDK1 inhibitor to neonatal mice. (E) Representative immunofluorescence images of ovarian sections from vehicle control (VC) and CDK1 inhibitor-treated (CDK1i) mice stained for MVH (green) and TAp63α (red). Square boxes indicate regions shown at higher magnification in the lower panels. Arrows indicate follicles expressing TAp63α. Scale bars: 100 μm (upper panels) and 10 μm (lower panels). (F) Quantification of total follicle numbers per ovary in vehicle control (VC) and CDK1 inhibitor-treated mice (n = 6 ovaries per group). Data are presented as mean ± SD. Statistical significance was determined using an unpaired two-tailed Student’s *t*-test. ns, not significant (P > 0.05).

### In Vivo Study (Neonatal mice)

To determine the role of CDK1 in ovarian reserve maintenance in vivo, neonatal mice were treated with a CDK1 inhibitor during a critical window for primordial follicle activation (7 to 11 dpp (days postpartum)). Mice received daily intraperitoneal injections of the inhibitor at a dose of 10 mg/kg (Fig. 3D). At the end of the treatment period, ovaries were collected, fixed, sectioned, and analyzed via immunofluorescence for MVH and TAp63α (Fig. 3E). Follicle counts revealed no significant differences between treated and control groups (Fig. 3F). To further assess the potential involvement of CDK1, the inhibitor dose was increased to 20 mg/kg, and the treatment regimen was repeated (Fig. 3D and E). However, even at this higher dose, no significant changes in follicle numbers were detected (Fig. 3F). These results suggest that CDK1 activity may not be essential for maintaining the ovarian follicle reserve, both in vitro and in vivo, under the experimental conditions employed.

### CDK1 is essential for maintaining oocyte viability under DNA damage induced by cisplatin

Proteomic analysis indicates CDK1 is upregulated in response to cisplatin, implying its involvement in the oocyte DNA damage response and repair mechanisms. Therefore, we administered 7 dpp mice with alternate-day treatments with a 10 mg/kg CDK1 inhibitor followed by 2 doses of 5 mg/kg of cisplatin via intraperitoneal injection up to 10 dpp (Fig. S3A). Ovaries were collected on the 10th day, fixed, sectioned, and subjected to immunofluorescence staining for MVH and TAp63α (Fig. S3B). Follicle quantification revealed no significant difference in follicle survival between the cisplatin-only and cisplatin + CDK1 inhibitor groups (Fig. S3C). Specifically, cisplatin-treated ovaries showed depletion of early-stage follicles, including primordial, primary, and secondary follicles, indicating that CDK1 inhibition did not further impact survival under these conditions.

To further investigate CDK1’s role in DNA repair, we reduced the cisplatin dose to a single 5 or 2.5 mg/kg injection, with and without CDK1 inhibition (Fig. S3A and B). Follicle quantification showed a 3.13-fold reduction in follicle numbers in the cisplatin-only group relative to controls, whereas the cisplatin + CDK1 inhibitor group exhibited a 7.18-fold reduction compared to controls and a 2.29-fold reduction compared to cisplatin alone (Fig. 4E). This indicates that CDK1 inhibition exacerbates follicle loss when DNA damage levels are moderate.

**Figure 4.**
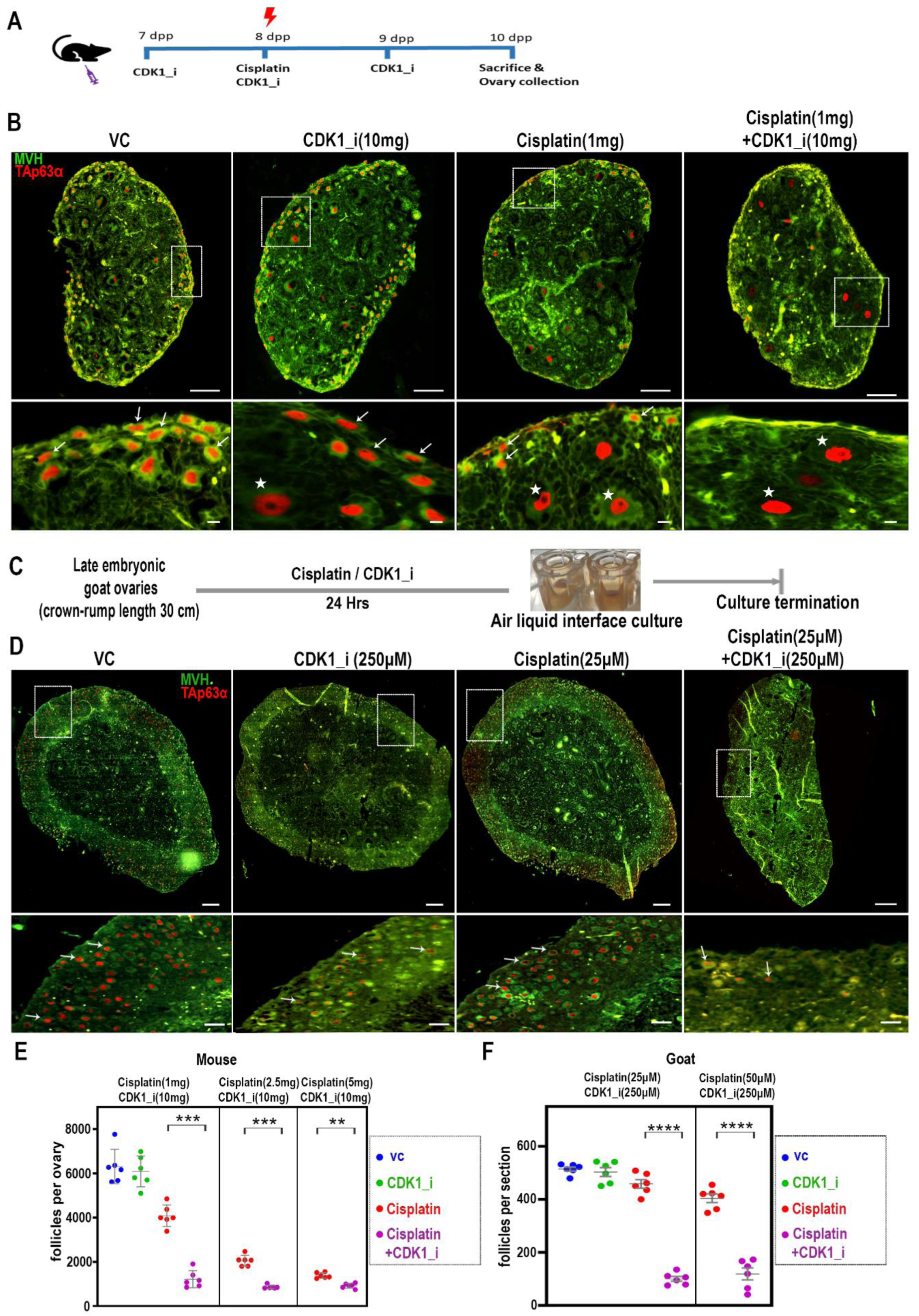
CDK1 is required for follicle survival following DNA damage. (A) Schematic overview of the in vivo experimental design. Neonatal mice were assigned to four treatment groups: Vehicle Control (VC), CDK1 inhibitor alone (CDK1i), Cisplatin (Cis), and Cisplatin plus CDK1 inhibitor (Cis + CDK1i). Treatments were administered by intraperitoneal injection. (B) Representative immunofluorescence images of ovarian sections from the indicated treatment groups stained for MVH (green) and TAp63α (red). White boxes indicate regions shown at higher magnification in the lower panels. Arrows indicate TAp63α-positive primordial follicles, and asterisks indicate TAp63α-positive secondary follicles. Scale bars: 100 μm (upper panels) and 10 μm (lower panels). (C) Schematic overview of the ex vivo experimental design using cultured late embryonic goat ovaries. Ovaries were treated with Vehicle Control (VC), CDK1 inhibitor alone (CDK1i), Cisplatin (Cis), or Cisplatin plus CDK1 inhibitor (Cis + CDK1i). (D) Representative immunofluorescence images of goat ovarian sections from the indicated treatment groups stained for MVH (green) and TAp63α (red). White boxes indicate regions shown at higher magnification in the lower panels. Arrows indicate TAp63α-positive follicles. Scale bars: 200 μm (upper panels) and 50 μm (lower panels). (E) Quantification of total follicle numbers per ovary in Vehicle Control (VC), CDK1 inhibitor (CDK1i), Cisplatin (Cis), and Cisplatin plus CDK1 inhibitor (Cis + CDK1i) mouse groups (n = 6 ovaries per group). (F) Quantification of total follicle numbers per section in Vehicle Control (VC), CDK1 inhibitor (CDK1i), Cisplatin (Cis), and Cisplatin plus CDK1 inhibitor (Cis + CDK1i) goat ovaries (n = 6 sections per group). Data are presented as mean ± SD. Statistical significance was determined using unpaired two-tailed Student’s *t*-tests for predefined pairwise comparisons between experimental groups. Significance levels are indicated as ****P ≤ 0.0001, ***P ≤ 0.001, and **P ≤ 0.01.

To examine CDK1’s role under minimal DNA damage, we administered a single 1 mg/kg cisplatin dose, with and without CDK1 inhibition (Fig. 4A and B). Follicle quantification showed a 1.57-fold reduction in cisplatin-only ovaries compared to the control group. In contrast, the cisplatin + CDK1 inhibitor group showed a 5.28-fold reduction compared to controls, underscoring CDK1’s critical role in oocyte survival under low-level DNA damage conditions (Fig. 4E and Fig. S3A-C).

To investigate the underlying mechanisms in higher-order vertebrates, we cultured goat ovaries and subjected them to the following treatments for up to 24 hours: cisplatin alone (25 or 50 µM), a CDK1 inhibitor alone, or a combination of cisplatin (25 or 50 µM) with the CDK1 inhibitor (Fig. 4C). Following treatment, ovarian tissues were fixed, sectioned, and immunostained for MVH and TAp63α to assess follicular integrity (Fig. 4D and Fig.S3D-E). Quantitative follicle analysis showed that CDK1 inhibition alone had no significant impact on follicle numbers. In contrast, treatment with cisplatin alone (25 or 50 µM) resulted in a modest 1.12-fold decrease in follicle count. Strikingly, the combination of cisplatin and CDK1 inhibition led to a 4.57-fold reduction in follicle numbers, indicating that CDK1 activity is essential for follicle survival in cisplatin-induced DNA damage (Fig. 4F). These findings indicate that CDK1-dependent repair mechanisms are conserved in higher-order vertebrates.

### CDK1 activation in oocyte DNA damage response and its regulation by Chk2

Despite ongoing oocyte turnover, the results suggest that CDK1 activity may not be essential for maintaining the ovarian reserve under normal conditions. However, it becomes crucial in response to moderate exogenous genotoxic stress. While CDK1 is primarily known for regulating the G2/M transition, it also plays a key role in the DNA damage response and homologous recombination repair. In oocytes arrested at prophase I, CDK1 remains inactive but can be further activated by DNA damage. In postnatal oocytes, DNA damage activates ATM and ATR kinases, leading to H2AX phosphorylation and Chk2 activation. Chk2 then phosphorylates TAp63α, initiating pathways for cell cycle arrest, repair, or apoptosis (model). Even in non-dividing cells such as oocytes, this Chk2-dependent checkpoint mechanism can partially activate CDK1 during exogenous DNA damage to maintain genomic integrity.

To investigate this hypothesis, 7 dpp mice were administered a single dose of Chk2 inhibitor (5 mg/kg) and CDK1 inhibitor (10 mg/kg) via intraperitoneal injection. On the 8th day, mice received a single dose of cisplatin (1 mg/kg) either with Chk2 inhibition alone or with combined Chk2 and CDK1 inhibition. On the 9th day, Chk2 and CDK1 inhibitors were administered again (Fig. 5A). The animals were sacrificed on the 10th day, and their ovaries were collected, fixed, sectioned, and subjected to immunofluorescence staining for MVH and TAp63α (Fig. 5B). Follicle quantification revealed that cisplatin alone shows a 1.29-fold follicle reduction. However, co-administration of cisplatin with CDK1 inhibition resulted in a further 4.66-fold reduction in follicle count. Cisplatin with Chk2 inhibition alone had slight protection, whereas the combination of cisplatin, Chk2, and CDK1 inhibitors produced a phenotype similar to cisplatin with Chk2 inhibition alone (Fig. 5C), suggesting that CDK1 activation may occur through the Chk2 checkpoint pathway.

**Figure 5.**
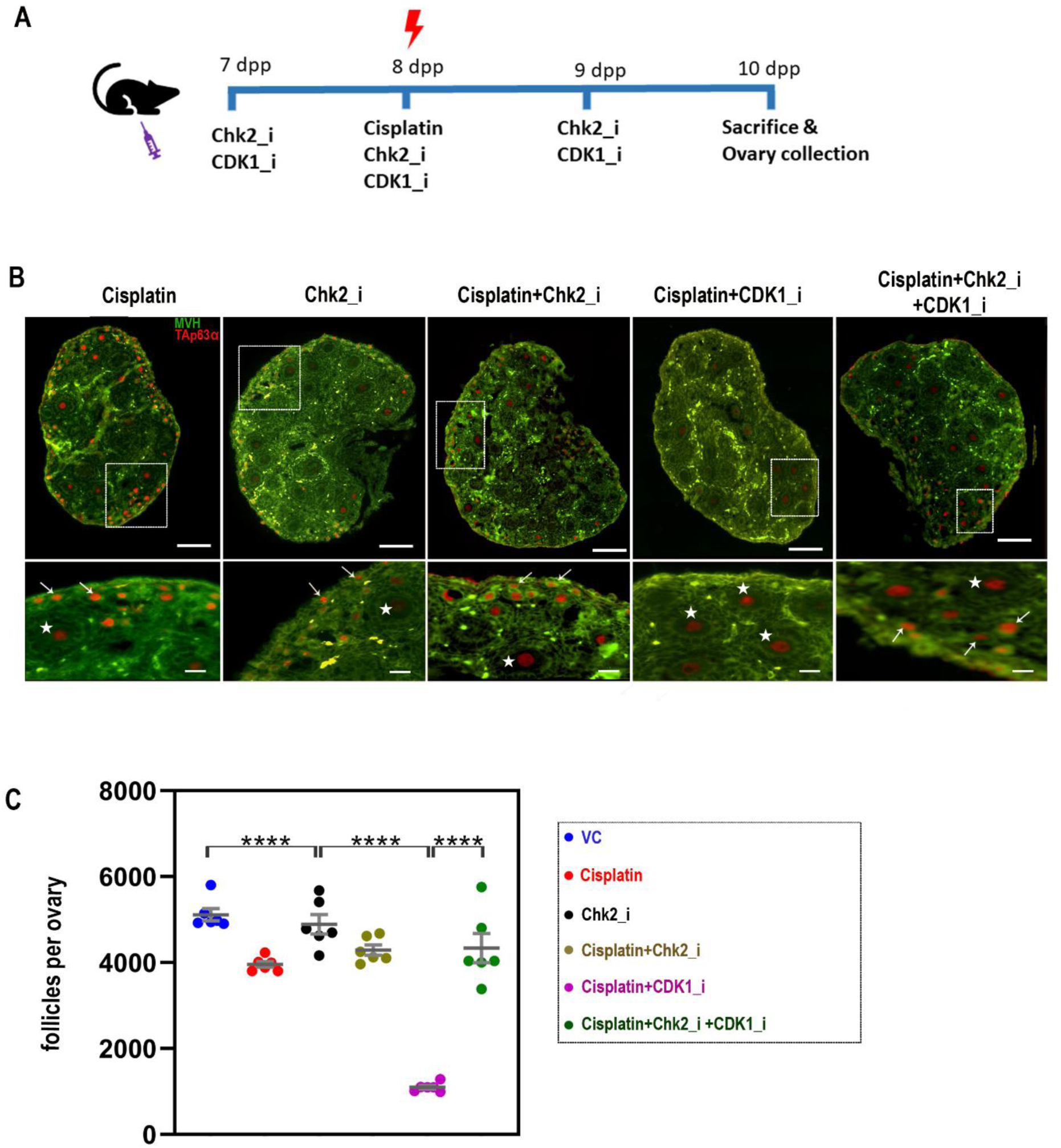
CDK1 activation is regulated by the DNA damage response through Chk2 signaling. (A) Schematic overview of the experimental design. Neonatal mice were assigned to four treatment groups: Vehicle Control (VC), Chk2 inhibitor alone (Chk2_i), Cisplatin (Cis), and Cisplatin plus Chk2 inhibitor (Cis + Chk2_i). Treatments were administered by intraperitoneal injection. (B) Representative immunofluorescence images of ovarian sections from the indicated treatment groups stained for MVH (green) and TAp63α (red). White boxes indicate regions shown at higher magnification in the lower panels. Arrows indicate TAp63α-positive primordial follicles, and asterisks indicate TAp63α-positive secondary follicles. Scale bars: 100 μm (upper panels) and 10 μm (lower panels). (C) Quantification of total follicle numbers per ovary in Vehicle Control (VC), Chk2 inhibitor (Chk2i), Cisplatin (Cis), and Cisplatin plus Chk2 inhibitor (Cis + Chk2i) groups (n = 6 ovaries per group). Data are presented as mean ± SD. Statistical significance was determined using unpaired two-tailed Student’s *t*-tests for predefined pairwise comparisons between experimental groups. Significance levels are indicated as ****P ≤ 0.0001.

### Chk2 regulates CDK1 activation through the WEE1–CDK1 checkpoint pathway during the oocyte DNA damage response

To investigate how Chk2 signaling influences CDK1 activity during DNA repair in dictyate-arrested oocytes, we examined the WEE1–CDK1 regulatory axis following cisplatin-induced genotoxic stress. Under physiological conditions, CDK1 activity is restrained by inhibitory phosphorylation at Thr14 and Tyr15, largely mediated by WEE1 family kinases, thereby maintaining meiotic arrest. We reasoned that activation of the DNA damage checkpoint might alter this regulatory network to facilitate repair-associated CDK1 activity (Fig. 6A). To test this hypothesis, neonatal mice were treated with cisplatin, Chk2 inhibitor, or a combination of cisplatin and Chk2 inhibitor as shown in the experimental scheme (Fig. 6B). Ovaries were subsequently analyzed by immunohistochemistry and western blotting to assess checkpoint activation and CDK1 regulatory components. Consistent with induction of DNA damage, cisplatin treatment resulted in a marked increase in γH2AX-positive oocytes compared with controls (2.09-fold, Fig. 6C–E and Fig.S5). This was accompanied by activation of the canonical oocyte DNA damage checkpoint, as evidenced by increased phospho-TAp63α and total TAp63α levels. Quantitative analysis showed that phospho-TAp63α increased by 1.1-fold relative to controls, confirming activation of the Chk2-dependent oocyte surveillance pathway (Fig. 6C, D, F&G and Fig.S5).

**Figure 6.**
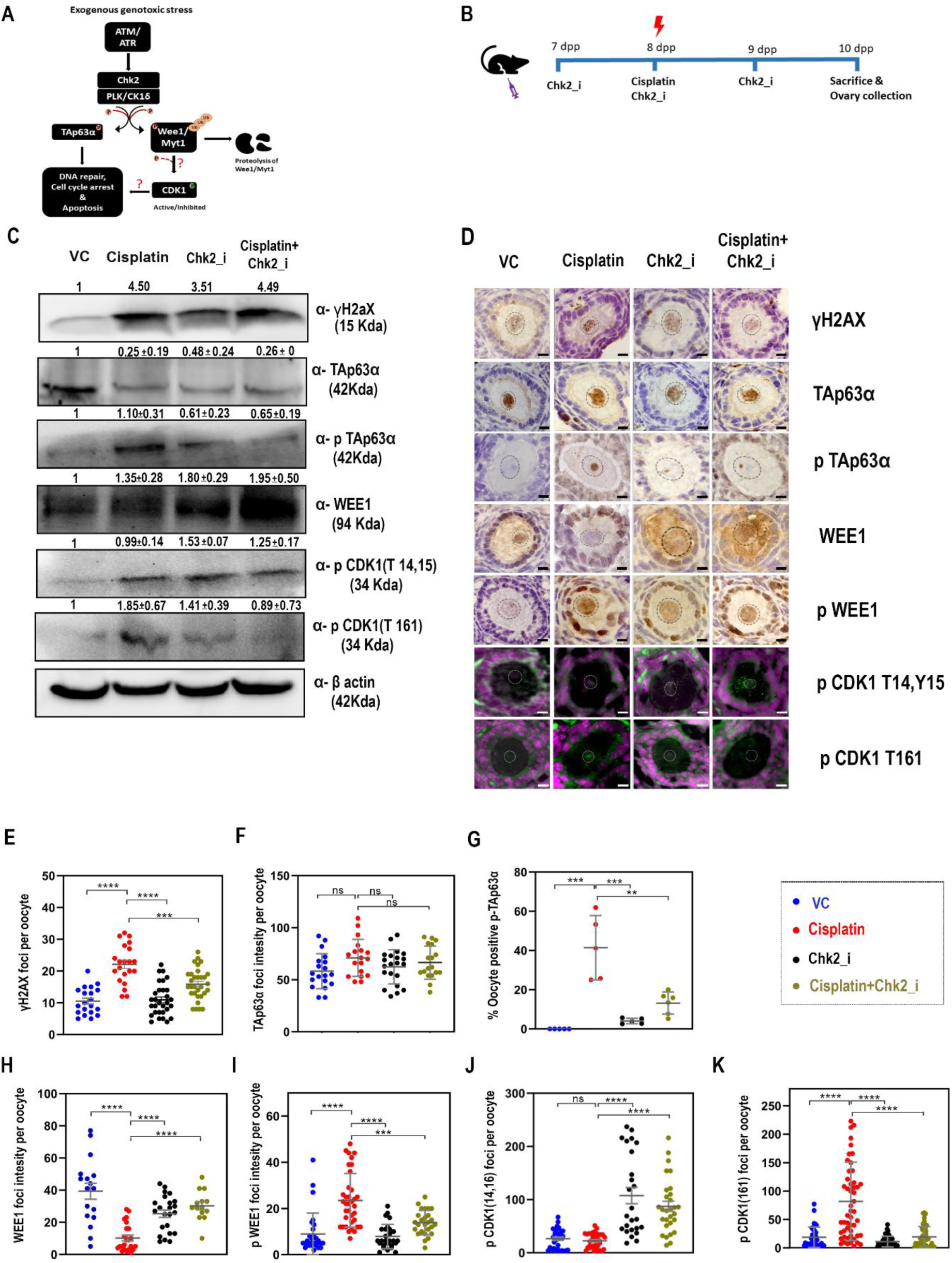
Chk2 regulates the WEE1–CDK1 checkpoint pathway during the oocyte DNA damage response. (A) Schematic model illustrating the proposed mechanism by which Chk2 regulates the WEE1–CDK1 checkpoint pathway in response to cisplatin-induced DNA damage in oocytes. (B) Experimental design. Neonatal mice were assigned to four treatment groups: Vehicle Control (VC), Chk2 inhibitor (Chk2_i), Cisplatin (Cis), and Cisplatin plus Chk2 inhibitor (Cis + Chk2_i). Treatments were administered by intraperitoneal injection, and ovaries were subsequently collected for molecular and histological analyses. As significant follicular changes were observed by 8-10 dpp, ovaries were collected at 9 dpp to evaluate early treatment-induced effects. (C) Representative western blots of ovarian lysates from the indicated treatment groups showing expression of γH2AX, TAp63α, phospho-TAp63α, WEE1, phospho-WEE1, phospho-CDK1 (T14/Y15), phospho-CDK1 (T161), and β-actin as a loading control. (D) Representative ovarian sections from each treatment group stained for γH2AX, TAp63α, phospho-TAp63α, WEE1, and phospho-WEE1 by immunohistochemistry. Sections were counterstained with hematoxylin. Immunofluorescence staining was performed for phospho-CDK1 (T14/Y15) and phospho-CDK1 (T161), with target proteins shown in green and nuclei counterstained with DAPI (magenta). Representative images depict secondary follicles. White and black circles indicate oocyte nuclei. (C&D) Representative western blot and immunohistochemistry (IHC) analysis of phospho-TAp63α using a mixture of antibodies against phospho-TAp63α (S160/162) and phospho-TAp63α (S455).Scale bar = 5 μm. (E) Quantification of γH2AX foci per oocyte. (F–I) Quantification of TAp63α, phospho-TAp63α, WEE1, and phospho-WEE1 expression in oocytes. TAp63α, WEE1, and phospho-WEE1 levels were quantified as staining intensity, whereas phospho-TAp63α was quantified as the percentage of positive oocyte. (J, K) Quantification of phospho-CDK1 (T14/Y15) and phospho-CDK1 (T161) foci per oocyte. Both staining intensity and foci were quantified in oocytes from primordial, primary, and secondary oocyte.Data are presented as mean ± SD. Statistical significance was determined using unpaired two-tailed Student’s *t*-tests for predefined pairwise comparisons between experimental groups. Significance levels are indicated as ****P ≤ 0.0001, ***P ≤ 0.001, **P ≤ 0.01, and *P ≤ 0.05.

We next examined whether activation of this checkpoint was associated with changes in the WEE1–CDK1 axis. Cisplatin exposure significantly reduced total WEE1 protein levels to 74% of control values while increasing phospho-WEE1 levels by 2.87-fold (Fig. 6C, D, H & I and Fig.S5). These changes were accompanied by a decrease in inhibitory CDK1 phosphorylation at Thr14/Tyr15 (14.89% reduction) and a corresponding increase in activating phosphorylation at Thr161 (4.5-fold increase) relative to controls, indicating a shift toward CDK1 activation during the DNA damage response.

To determine whether these molecular changes were dependent on Chk2 activity, ovaries were co-treated with cisplatin and Chk2 inhibitor. Although γH2AX levels remained elevated, indicating the continued presence of DNA lesions, inhibition of Chk2 substantially attenuated downstream checkpoint activation. Both phospho-TAp63α and total TAp63α levels were reduced to 67.89% and 6.3% of those observed in cisplatin-treated ovaries, respectively (Fig. 6C, D, F&G and Fig.S5). Notably, Chk2 inhibition restored WEE1 expression more than control levels and normalized phospho-WEE1 abundance (68.90% and 4.8%, respectively) (Fig. 6C, D, H&I and Fig.S5). Restoration of WEE1 was accompanied by recovery of inhibitory CDK1 phosphorylation at Thr14/Tyr15 (4.89-fold increase compared with cisplatin alone) and a significant reduction in activating CDK1 phosphorylation at Thr161 (86.49% decrease; (Fig. 6C, D, J&K and Fig.S5). These findings demonstrate that the DNA damage-induced shift in CDK1 phosphorylation status is dependent on Chk2 signaling. Collectively, these results identify Chk2 as an upstream regulator of the WEE1–CDK1 checkpoint pathway in damaged dictyate-arrested oocytes. DNA damage-induced Chk2 activation is associated with modulation of WEE1, reduced inhibitory phosphorylation of CDK1, and increased activating phosphorylation of CDK1, whereas pharmacological inhibition of Chk2 reverses these molecular changes. Together with the follicle survival data presented in Figure 5, these findings position CDK1 downstream of Chk2 signaling during the oocyte response to genotoxic stress.

### CDK1 inhibition suppresses homologous recombination and is associated with increased expression of NHEJ and NER markers following oocyte DNA damage

The preceding experiments demonstrated that CDK1 activity is required for maintaining follicle survival during cisplatin-induced genotoxic stress and that inhibition of CDK1 results in persistent DNA damage accumulation. Because homologous recombination (HR) is the predominant pathway for repairing DNA double-strand breaks in oocytes, we next investigated whether CDK1 influences HR-associated repair responses through regulation of RAD51 activation and recruitment. We further examined whether suppression of HR was accompanied by changes in alternative DNA repair pathways (Fig. 7A).

**Figure 7.**
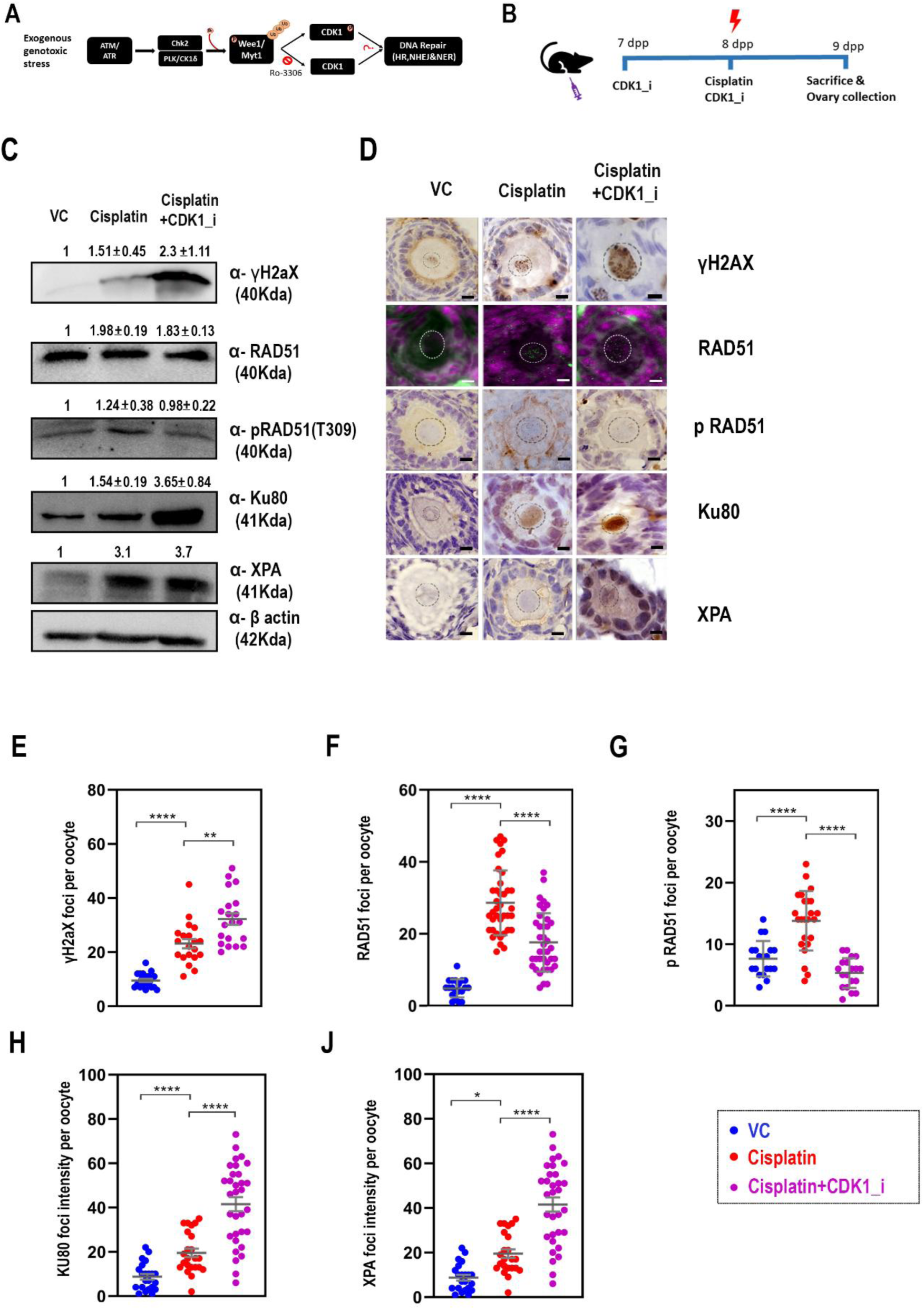
CDK1 inhibition suppresses homologous recombination and promotes the expression of NHEJ and NER repair markers following oocyte DNA damage. (A) Schematic model illustrating the proposed role of CDK1 in regulating DNA repair pathway choice in oocytes following cisplatin-induced DNA damage. Active CDK1 promotes RAD51-dependent homologous recombination (HR), whereas CDK1 inhibition suppresses HR-associated repair and is associated with increased expression of non-homologous end joining (NHEJ) and nucleotide excision repair (NER) pathway markers. (B) Experimental design. Neonatal mice were assigned to three treatment groups: Vehicle Control (VC), Cisplatin (Cis), and Cisplatin plus CDK1 inhibitor (Cis + CDK1_i). As significant follicular changes were observed by 8-10 dpp, ovaries were collected at 9 dpp to evaluate early treatment-induced effects. Treatments were administered by intraperitoneal injection, and ovaries were subsequently collected for molecular and histological analyses. (C) Representative western blot analysis of ovarian lysates from the indicated treatment groups showing expression of γH2AX, RAD51, phospho-RAD51, KU80, XPA, and β-actin as a loading control. (D) Representative ovarian sections from each treatment group stained for γH2AX, KU80, and XPA by immunohistochemistry. Sections were counterstained with hematoxylin. Immunofluorescence staining was performed for RAD51 and phospho-RAD51, with target proteins shown in green and nuclei counterstained with DAPI (magenta). Representative images depict secondary follicles. White and black circles indicate oocyte nuclei. Scale bar = 5 μm. (E) Quantification of γH2AX foci per oocyte. (F) Quantification of RAD51 and phospho-RAD51 foci per oocyte. (G, H) Quantification of KU80 and XPA staining intensities in oocytes from primordial, primary, and secondary oocyte. Data are presented as mean ± SD. Statistical significance was determined using unpaired two-tailed Student’s *t*-tests for predefined pairwise comparisons between experimental groups. Significance levels are indicated as ****P ≤ 0.0001, ***P ≤ 0.001, **P ≤ 0.01, and *P ≤ 0.05.

To address this question, neonatal mice were treated with cisplatin alone or with cisplatin and CDK1 inhibitor according to the experimental design shown in Fig. 7B. Ovaries were collected at 9 dpp and analyzed by immunohistochemistry, immunofluorescence, and western blotting. As expected, cisplatin treatment induced a robust DNA damage response, as evidenced by increased γH2AX staining in primordial, primary, and secondary follicles (2.42-fold versus control, Fig. 7C–E and Fig.S7). Under these conditions, RAD51 accumulation was increased within oocytes, consistent with activation of HR-associated repair processes. Quantitative analysis revealed a 5.83-fold increase in RAD51 signal intensity relative to controls. In parallel, phospho-RAD51 levels were elevated following cisplatin exposure (1.81-fold increase,), indicating enhanced RAD51 activation during the DNA damage response.

To determine whether RAD51 activation was dependent on CDK1 activity, ovaries were co-treated with cisplatin and CDK1 inhibitor. Despite persistent γH2AX accumulation, RAD51 localization was reduced across all follicle classes compared with cisplatin treatment alone (37.07% reduction; Fig. 7C, F and Fig.S7). Similarly, phospho-RAD51 staining was significantly decreased following CDK1 inhibition (61.5% reduction). These observations suggest that CDK1 activity contributes to efficient RAD51 activation and recruitment in DNA-damaged oocytes.

Western blot analyses corroborated the immunostaining results. Cisplatin treatment increased both total RAD51 and phospho-RAD51 protein levels by 1.9-fold and 1.24-fold, respectively, relative to controls (Fig. 7C, D). In contrast, CDK1 inhibition reduced RAD51 and phospho-RAD51 abundance to 9.01% and 14% of cisplatin-treated levels, respectively. Notably, γH2AX levels remained elevated in the cisplatin plus CDK1 inhibitor group (2.30-fold versus control), indicating persistence of unrepaired DNA lesions despite activation of the DNA damage response.

Because compromised HR repair can lead to increased reliance on alternative repair mechanisms, we next examined markers associated with non-homologous end joining (NHEJ) and nucleotide excision repair (NER). Cisplatin treatment induced moderate increases in the NHEJ factor Ku80 and the NER factor XPA. However, both proteins were further elevated following CDK1 inhibition. Immunohistochemical analysis revealed a 2.23-fold increase in Ku80-positive oocytes and a 1.57-fold increase in XPA-positive oocytes relative to cisplatin treatment alone (Fig. 7C, D, H&J and Fig.S7). Consistent with these findings, western blot analysis showed that Ku80 and XPA protein levels increased by 3.6-fold and 3.7-fold, respectively, in the cisplatin plus CDK1 inhibitor group compared with cisplatin-treated ovaries. Collectively, these findings indicate that inhibition of CDK1 attenuates RAD51 activation and recruitment in response to DNA damage, resulting in persistent γH2AX accumulation and incomplete lesion resolution. This impairment is accompanied by increased expression of NHEJ- and NER-associated repair factors, suggesting compensatory engagement of alternative DNA repair pathways when HR-associated repair responses are compromised. Together, these data support a role for CDK1 in facilitating efficient homologous recombination-associated DNA repair in dictyate-arrested oocytes exposed to genotoxic stress.

### CDK1 inhibition does not induce premature meiotic resumption in dictyate-arrested oocytes

Because CDK1 is a well-established regulator of meiotic resumption, we investigated whether the follicle survival phenotype observed following CDK1 inhibition could be attributed to alterations in oocyte cell-cycle status rather than a direct effect on DNA repair. To address this possibility, we assessed meiotic status following CDK1 inhibition using the experimental strategy outlined in Fig. 8A.

**Figure 8.**
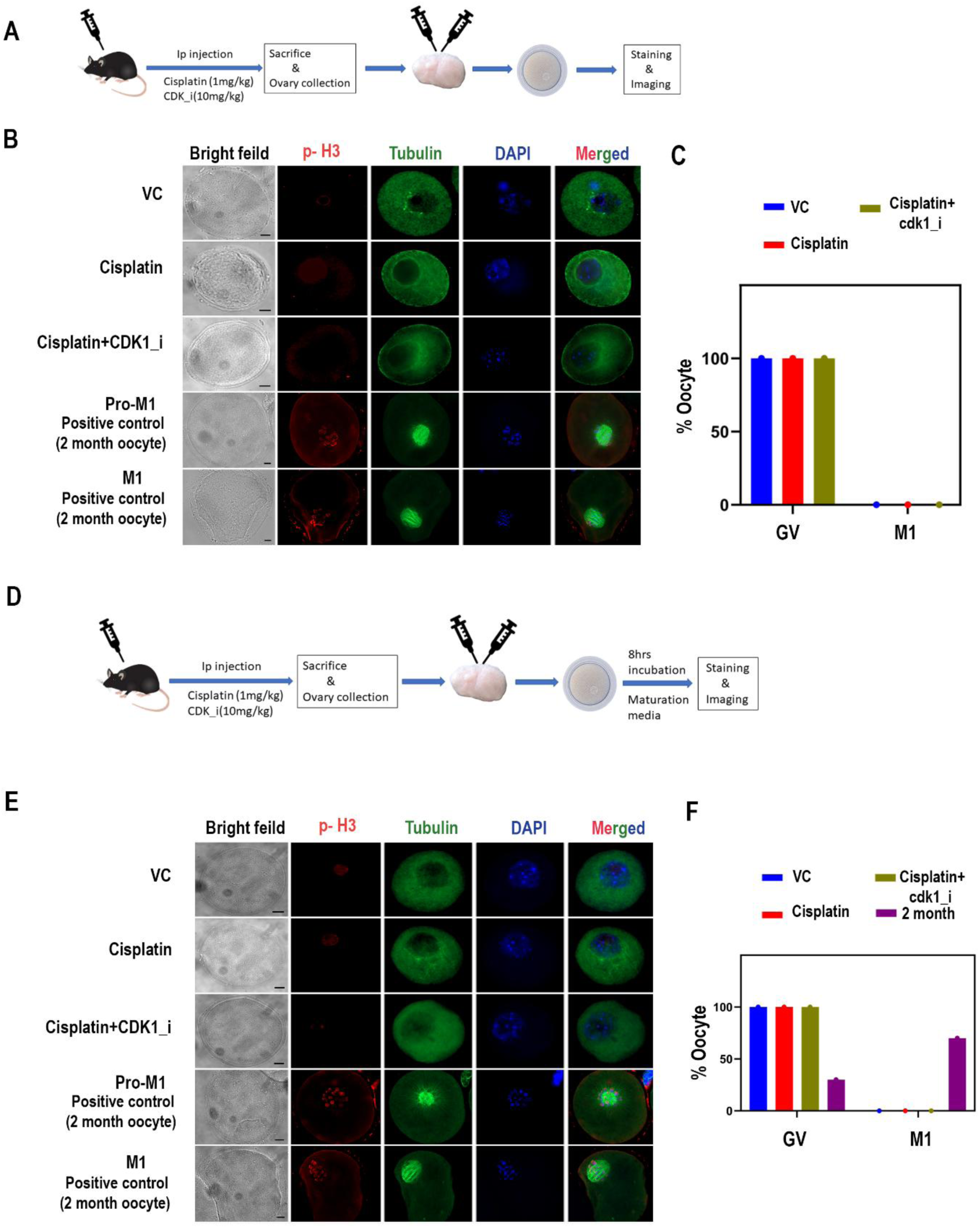
DNA damage-associated CDK1 activation is insufficient to drive meiotic maturation in oocytes. (A) Schematic overview of the experimental design. Neonatal mice were assigned to three treatment groups: Vehicle Control (VC), Cisplatin (Cis), and Cisplatin plus CDK1 inhibitor (Cis+CDK1_i). As significant follicular changes were observed by 8-10 dpp, ovaries were collected at 9 dpp to evaluate early treatment-induced effects. Mice were sacrificed at 9 dpp. Treatments were administered by intraperitoneal injection, and ovaries were subsequently collected for molecular and histological analyses. (B) Representative immunofluorescence images of oocytes from the indicated treatment groups stained for phospho-histone H3 (pH3), α-tubulin, and DNA (DAPI). Individual fluorescence channels and merged images are shown. Scale bar = 10 μm. (C) Quantification of oocytes according to meiotic stage and maturation status, including germinal vesicle (GV), and metaphase I (MI) stage oocytes. (D) Schematic overview of the in vitro maturation (IVM) experiment. Following the indicated in vivo treatments (VC, Cis, and Cis + CDK1_i), oocytes were isolated and cultured in vitro for 8 h prior to analysis. (E) Representative immunofluorescence images of in vitro matured oocytes from the indicated treatment groups stained for phospho-histone H3 (pH3), α-tubulin, and DNA (DAPI). Individual fluorescence channels and merged images are shown. Scale bar = 10 μm. (F) Quantification of oocyte maturation status following in vitro culture, showing the proportion of oocytes at the GV and MI stages. Data are presented as mean ± SD. Statistical significance was determined using unpaired two-tailed Student’s *t*-tests for predefined pairwise comparisons between experimental groups. Significance levels are indicated as ****P ≤ 0.0001, ***P ≤ 0.001, **P ≤ 0.01, and *P ≤ 0.05.

Two complementary approaches were employed. In the first, prepubertal mice were treated in vivo with cisplatin alone or cisplatin together with a CDK1 inhibitor, after which oocytes were isolated and analyzed (Fig. 8A). In the second, isolated oocytes were cultured for an additional 8 h under in vitro maturation conditions to evaluate their capacity to undergo meiotic resumption (Fig. 8D). To validate the detection of meiotic progression markers, fully grown metaphase I (MI)-stage oocytes obtained from 2-month-old mice were included as positive controls. Oocytes were stained for phospho-histone H3 (Ser10), a marker of chromosome condensation, together with α-tubulin to visualize spindle assembly and chromosome organization (Fig. 8B). As expected, MI-stage oocytes displayed strong phospho-histone H3 staining and well-organized bipolar spindles, confirming successful meiotic progression and validating the staining approach. In contrast, dictyate-arrested oocytes isolated from cisplatin-treated ovaries lacked phospho-histone H3 staining and showed no evidence of spindle assembly. Importantly, oocytes isolated from ovaries treated with cisplatin and CDK1 inhibitor exhibited a similar phenotype, remaining morphologically arrested and devoid of chromosome condensation or spindle structures characteristic of meiotic resumption (Fig. 8B). To further evaluate meiotic competence, isolated oocytes were cultured under in vitro maturation conditions for 8 h. Consistent with the in vivo observations, oocytes exposed to CDK1 inhibition failed to undergo efficient meiotic progression, with no oocytes reaching metaphase I compared with control oocytes (100% versus 70%, Fig. 8F). These findings confirm neither direct examination of oocytes following in vivo treatment nor subsequent in vitro maturation assays provided evidence that CDK1 inhibition induced premature meiotic resumption in dictyate-arrested oocytes. Therefore, the increased DNA damage accumulation, apoptosis, and follicle depletion observed following CDK1 inhibition are unlikely to result from aberrant activation of the meiotic program and instead support a direct role for CDK1 in the oocyte DNA damage response.

## Discussion

Maintenance of genomic integrity in dictyate-arrested oocytes is essential for preserving female fertility throughout reproductive life. Because primordial follicle oocytes remain arrested in prophase I for prolonged periods, they are continuously exposed to endogenous and environmental sources of DNA damage. Although checkpoint-mediated elimination of severely damaged oocytes has been extensively studied, the mechanisms that allow dormant oocytes to actively repair DNA lesions while maintaining meiotic arrest remain poorly understood. In the present study, we identify a previously unrecognized Chk2–CDK1 signaling axis that promotes homologous recombination (HR)-mediated DNA repair in postnatal oocytes exposed to genotoxic stress. Our findings demonstrate that CDK1, beyond its canonical role in meiotic progression, functions as a repair-associated kinase that supports RAD51-dependent DNA repair and preserves follicle survival under moderate DNA damage conditions.

Proteomic analysis of cisplatin-treated goat ovaries revealed significant enrichment of proteins associated with DNA damage signaling, cell-cycle regulation, and repair pathways, with CDK1 emerging as one of the most prominently upregulated kinases. This observation was accompanied by increased γH2AX accumulation and activation of Chk2, confirming robust induction of the DNA damage response. Although CDK1 is classically associated with the G2/M transition and meiotic resumption, increasing evidence from somatic systems suggests that CDK1 also regulates homologous recombination through phosphorylation of DNA repair proteins. Our findings extend this concept to dictyate-arrested oocytes and suggest that transient CDK1 activation can occur independently of meiotic progression to facilitate genome maintenance.

A key finding of this study is that CDK1 signalling is dispensable for maintaining the ovarian follicle pool under physiological conditions but becomes essential during genotoxic stress. Pharmacological inhibition of CDK1 alone did not significantly affect follicle numbers in either goat ovarian cultures or neonatal mouse ovaries, indicating that basal follicle survival does not require sustained CDK1 activity during meiotic arrest. In contrast, CDK1 inhibition markedly exacerbated follicle loss following cisplatin exposure, particularly under moderate and low-dose DNA damage conditions. These observations suggest that dormant oocytes possess a threshold-dependent repair response in which CDK1-dependent repair mechanisms become critically important when DNA damage remains potentially repairable. Under excessive damage, however, apoptotic pathways likely dominate irrespective of CDK1 activity.

Mechanistically, our data indicate that CDK1 promotes homologous recombination-associated repair through regulation of RAD51 phosphorylation and accumulation at DNA damage foci. RAD51 is indispensable for homology search and strand invasion during HR-mediated repair of DNA double-strand breaks. We observed robust RAD51 recruitment and increased phospho-RAD51 levels following cisplatin treatment, whereas CDK1 inhibition markedly reduced RAD51 localization and phosphorylation despite persistent γH2AX accumulation. These findings suggest that CDK1 activity is required for efficient RAD51 loading onto damaged chromatin in oocytes. Previous studies in yeast demonstrated that Cdc28-dependent phosphorylation enhances Rad51 DNA-binding activity during homologous recombination. Our results suggest that a similar evolutionarily conserved mechanism may operate in mammalian oocytes to maintain genome integrity during prolonged meiotic arrest.

An important conceptual question arising from this work is how CDK1 can participate in DNA repair without triggering meiotic resumption. In dictyate-arrested oocytes, CDK1 activity is normally suppressed by inhibitory phosphorylation at Thr14 and Tyr15 through the actions of WEE1 and MYT1, thereby preventing premature meiotic entry. Our findings suggest that DNA damage induces a transient and partial activation of CDK1 rather than the global activation associated with meiotic progression. Notably, CDK1 activation under DNA damage conditions did not induce meiotic resumption, as oocytes retained dictyate-stage morphology and lacked phospho-histone H3 signals associated with chromosome condensation. These observations support the possibility that DNA damage induces a spatially or temporally restricted pool of active CDK1 dedicated to repair-associated signaling rather than meiotic entry. More broadly, these findings challenge the traditional view that CDK1 activity in oocytes is exclusively associated with meiotic resumption. Instead, our data support the existence of context-dependent CDK1 signaling, whereby limited activation of CDK1 can be uncoupled from cell-cycle progression and redirected toward genome maintenance functions. Such compartmentalized or threshold-dependent kinase activity may represent a general strategy by which long-lived quiescent cells utilize cell-cycle regulators to preserve genomic integrity while maintaining a non-dividing state.

Our data further identify Chk2 as an upstream regulator linking DNA damage sensing to CDK1 activation. Chk2 is well established as a critical checkpoint kinase in oocytes, where it phosphorylates TAp63α and promotes either DNA repair or apoptotic elimination depending on the severity of damage. Here, we demonstrate that inhibition of Chk2 abolished CDK1 activation and reversed the molecular changes associated with DNA damage-induced modulation of the WEE1–CDK1 axis. Cisplatin treatment reduced WEE1 abundance and inhibitory phosphorylation of CDK1 while increasing activating phosphorylation at Thr161, consistent with transient CDK1 activation during DNA repair. In contrast, Chk2 inhibition restored WEE1 levels and inhibitory CDK1 phosphorylation, suggesting that Chk2 signaling is required to relieve WEE1-mediated suppression of CDK1 during the oocyte DNA damage response.

The potential involvement of a Chk2–CK1–WEE1 signaling cascade is particularly intriguing. Previous studies have demonstrated that Chk2 primes TAp63α for subsequent CK1-mediated phosphorylation during oocyte quality control, whereas studies in mitotic systems have shown that CK1-dependent phosphorylation can contribute to WEE1 destabilization and ubiquitin-mediated degradation. Consistent with these observations, we found that CK1α expression increased following cisplatin-induced DNA damage and was reduced upon Chk2 inhibition, supporting its placement downstream of Chk2 signalling during the oocyte DNA damage response (Fig. S6). These findings raise the possibility that CK1α may contribute to modulation of the WEE1–CDK1 regulatory axis in damaged oocytes. Such a mechanism could facilitate a transient DNA damage-associated CDK1 activation signature while preserving meiotic arrest, thereby enabling activation of homologous recombination-associated repair pathways without triggering meiotic resumption. Although additional biochemical studies will be required to directly determine whether CK1α regulates WEE1 stability or CDK1 activity in oocytes, our findings provide a framework linking checkpoint signalling, CK1α induction, and repair-associated CDK1 regulation during the oocyte DNA damage response.

Another important implication of this study is the observation that suppression of CDK1-dependent homologous recombination is associated with increased expression of alternative DNA repair pathway markers. CDK1 inhibition caused persistent γH2AX accumulation together with increased Ku80 and XPA expression, which may indicate the enhanced activation of non-homologous end joining (NHEJ) and nucleotide excision repair (NER) pathways. While these pathways may partially compensate for defective HR repair, they are generally considered less accurate for repairing DNA double-strand breaks in meiotic cells. Increased reliance on NHEJ may therefore promote mutagenic repair and compromise oocyte genomic stability. This finding may have important implications for reproductive aging and chemotherapy-induced infertility, where repeated genotoxic stress could progressively impair HR efficiency and alter the expression of DNA repair pathway-associated factors.

The translational implications of these findings are substantial. Fertility preservation remains a major challenge for young cancer patients undergoing chemotherapy, and current strategies primarily focus on ovarian tissue cryopreservation or suppression of follicle activation. Our findings suggest an alternative approach centered on enhancing endogenous DNA repair pathways within dormant oocytes. Modulation of CDK1 activity or stabilization of RAD51-dependent HR repair machinery may represent potential strategies to improve oocyte survival during chemotherapy. Moreover, controlled manipulation of the Chk2–CDK1 axis may provide a means to balance DNA repair and apoptotic elimination within the ovarian reserve, thereby preserving long-term reproductive function. The present findings may also have implications for natural reproductive aging. Accumulation of unrepaired DNA damage is a hallmark of aging oocytes, and age-associated declines in homologous recombination factors, including RAD51 and BRCA-associated proteins, have been reported. A progressive reduction in CDK1-dependent repair signaling could therefore contribute to the diminished DNA repair capacity observed in aging ovarian reserves, ultimately accelerating follicle attrition and reproductive decline.

An important strength of this study is the use of both goat and mouse models to investigate DNA repair in dictyate-arrested oocytes. Fetal goat ovaries contain a large population of dictyate-stage oocytes and share several developmental features with the human ovary, providing a physiologically relevant system for studying mechanisms that preserve the ovarian reserve. The conservation of DNA damage-induced CDK1 activation and repair responses across these species supports the broader relevance of this pathway in mammalian oocyte biology.

Despite these advances, several questions remain. Although our findings demonstrate that CDK1 activity is required for efficient RAD51 phosphorylation and recruitment during the oocyte DNA damage response, the direct molecular mechanisms linking CDK1 to the homologous recombination machinery remain to be defined. Likewise, while our data support a model in which Chk2 signaling regulates the WEE1–CDK1 axis, the proposed involvement of CK1-dependent WEE1 destabilization will require further biochemical validation.

In addition, the present study primarily focused on repair responses within oocytes, although several DNA damage response proteins were also detected in surrounding granulosa and pre-granulosa cells. Emerging evidence suggests that bidirectional communication between germ cells and the somatic follicular compartment contributes to both follicle maintenance and responses to genotoxic stress. While the most pronounced changes in γH2AX, phospho-TAp63α, RAD51, and phospho-RAD51 were observed within oocytes and closely correlated with follicle survival, potential contributions from somatic cells cannot be excluded. Future studies employing cell type-specific genetic approaches will be important for defining how oocyte- and granulosa cell-derived signals cooperate to coordinate DNA repair and follicle preservation.

In summary, our study has unveiled a previously unrecognized role for CDK1 in maintaining genome integrity in dictyate-arrested oocytes. We propose a model in which DNA damage activates Chk2 signaling, leading to modulation of the WEE1–CDK1 inhibitory axis and transient CDK1 activation. Activated CDK1 subsequently promotes RAD51 phosphorylation and accumulation at DNA damage foci, supporting homologous recombination-associated repair, thereby preserving follicle survival under moderate genotoxic stress. Disruption of this pathway impairs HR repair and shifts oocytes toward less efficient repair mechanisms, ultimately accelerating follicle loss. Together, these findings redefine CDK1 as a repair-associated kinase in the ovarian reserve and provide new insight into how dormant oocytes balance genome maintenance with long-term reproductive survival.

## Material and methods

### Animal Experiments

Female C57BL/6 mice were used for in vivo studies and housed under standard laboratory conditions with a 12-hour light/dark cycle. All animal procedures were conducted in accordance with the guidelines approved by the Institutional Animal Ethics Committee of the National Institute of Animal Biotechnology (IAEC/2024/NIAB/17/HBDPR). Mice received intraperitoneal (i.p.) injections of the following agents at concentrations specified in the corresponding figure legends: Cisplatin (HY-17394, MedChemExpress) as a DNA-damaging agent, BML-277 (HY-13946, MedChemExpress) as a Chk2 inhibitor, and Ro-3306 (S7747, Selleckchem) as a CDK1 inhibitor.

### Sample Collection

Fetal ovaries were obtained from Osmanabadi goat breed (Capra hircus) fetuses collected from local slaughterhouses. Fetuses were selected based on a crown-rump length of approximately 30 cm, indicative of comparable developmental stages. Ovaries were dissected under sterile conditions and washed 4–5 times in TCM199 medium supplemented with antibiotics (penicillin-streptomycin) to minimize microbial contamination. Only ovaries of similar size and morphology were selected for use in triplicate biological replicates.

### Air-Liquid Interface Culture System

Fetal ovaries were cultured using an air-liquid interface system to support tissue viability and development. Dissected ovaries were placed on a sterile porous membrane or support structure, ensuring contact with the culture medium from below and exposure to air from above. Cultures were maintained at 37°C in a humidified incubator with 5% CO₂. The culture medium consisted of TCM199 supplemented with 3% bovine serum albumin (BSA), 10% goat follicular fluid, and 1X penicillin-streptomycin. Medium was replenished regularly, and tissue viability was monitored throughout the culture period.

### Induction of DNA Damage

To induce DNA damage in cultured ovaries, cisplatin was added to the culture medium at a final concentration of 100 µM. Ovaries were incubated with cisplatin for 24 hours under standard culture conditions. Following treatment, ovaries were immediately snap-frozen in liquid nitrogen and stored at –80°C until further processing.

### Mass spectroscopy

Fetal ovaries from vehicle control and DNA damage-induced conditions were lysed in lysis buffer containing 1% sodium deoxycholate in 50 mM ammonium bicarbonate. Protein extraction and sample preparation were performed as previously described by Kumar et al. (2024)(34). Briefly, lysates were reduced with dithiothreitol at 57°C for 1 hour, followed by alkylation with iodoacetamide in the dark at room temperature for 1 hour. Proteins were digested overnight (14 hours) at 37°C using trypsin at an enzyme-to-protein ratio of 1:30. After digestion, SDC was precipitated by adding formic acid to a final concentration of 20%. The resulting peptide mixtures were filtered through a 10 kDa molecular weight cut-off filter (Thermo Fisher Scientific) to remove undigested proteins. Salts and other contaminants were removed using C18 desalting columns (Thermo Fisher Scientific), and the eluted peptides were vacuum-dried. Dried peptides were reconstituted in 0.1% formic acid, and 1–2 µg of peptides per sample were injected for LC-MS/MS analysis. Peptides were separated over a 180-minute gradient and analyzed using a Thermo Fisher Scientific mass spectrometer. Data acquisition was followed by processing in the Proteome Discoverer software.

### Proteome Analysis

Proteomic data analysis was performed as described by Kumar et al. (2024) (34). Protein identification was carried out using Proteome Discoverer version 2.5 (Thermo Fisher Scientific), with searches against the UniProtKB databases for goat (UP000291000) and sheep (UP000002356). Label-free quantification (LFQ) was used to determine relative protein abundance across vehicle control and DNA damage-induced samples. Protein intensities were normalized, and differential expression was considered significant with a log2 fold change >1 or <–1 and a p-value <0.05. Venn diagrams were used to visualize the overlap in protein identifications across conditions. Principal Component Analysis plots illustrated variation within and between sample groups. Heat maps were generated to depict differentially regulated proteins between the two conditions. Gene ontology enrichment analysis was performed using an over-representation test in FunRich v3.1.3. Volcano plots were used to highlight significantly upregulated and downregulated proteins. PCA and heat maps were generated using the ClustVis online tool.

### Data availability

The mass spectrometry proteomics data have been deposited in the Mass Spectrometry Interactive Virtual Environment (Massive) repository with the dataset identifier MSV000097779.

### Ovary fixation and sectioning

Ovary samples were processed as previously published by Kumar et al. (2024) with some modifications(34). Ovaries were fixed in 10% formalin for 24 hrs based on the size of the ovary, then incubated with sucrose gradient with 10%, 20%, and 30% series for 24 hrs each and processed for cryotome. Further, thin 5 µm sections were used for immunostaining.

### Immunofluorescence

Ovarian tissue sections were subjected to antigen retrieval by boiling in antigen retrieval buffer at 95°C for 50 minutes, followed by cooling at room temperature for 20 minutes. The sections were then blocked with antibody dilution buffer (ADB) twice for 15 minutes each to reduce nonspecific binding. After blocking, primary antibodies: mouse anti-p63 (CM163B, 1:500), rabbit anti-MVH (ab13840, 1:500), rabbit anti-phospho-CDK1 (T14, Y15) (44-686G; 1:100), rabbit anti-phospho-CDK1 (T161) (PA5-117191; 1:100) and rabbit anti-Rad51 (PC130, 1:250), and rabbit anti- phospho-Rad51(T309) (BS-5677R; 1:100), were applied to the sections and incubated overnight at humid chamber. The following day, slides were washed thrice with TBST (Tris-buffered saline with Tween-20) for 5 minutes per wash. Slides were then incubated for 1 hour at 37°C in the dark with fluorophore-conjugated secondary antibodies: goat anti-rabbit Alexa Fluor 488 (A32731, 1:2000) and goat anti-mouse Alexa Fluor 555 (A16076, 1:2000). After incubation, slides were again washed three times with TBST and counterstained with DAPI for 2 minutes to visualize nuclei. Finally, the sections were mounted using DPX mounting medium, and images were captured using a fluorescence microscope.

### Immunohistochemistry

For immunohistochemical analysis, 5 µm ovary sections were deparaffinized, rehydrated, and subjected to antigen retrieval by heating in antigen retrieval buffer at 95°C for 50 minutes, then cooled to room temperature for 20 minutes. Sections were blocked with antibody dilution buffer (ADB) twice for 15 minutes at room temperature. After blocking, the sections were incubated overnight at humid chamber with the primary antibodies: mouse anti phospho-γH2A.X (S 139) (05-636; 1:250), rabbit anti-WEE1(PA5-29303; 1:200), rabbit anti-phospho-WEE1(S 139) (PA5-105559; 1:200), mouse anti-p63 (CM163B; 1:500),rabbit anti-p63(S160/162) (4981S; 1:50), rabbit anti-p63(S 455) (STJ90859; 1:50),rabbit anti-KU80(2180; 1:500), and rabbit anti-XPA (PA5-119824; 1:200). After incubation, slides were washed three times with TBST for 5 minutes each. A biotin-labeled goat anti-mouse secondary antibody (A16076, 1:1000) and goat anti-rabbit secondary antibody (B-2770, 1:1000) were applied and incubated for 1 h at 37°C.. The slides were again washed three times with TBST and incubated with the Vectastain ABC reagent (PK-6100) for 1 hour. After a brief wash with PBST for 5 minutes, DAB substrate (SK-4100) was applied for 90 minutes to develop color. Finally, sections were counterstained with hematoxylin (40362), dehydrated, mounted using DPX, and visualized under a bright-field microscope.

### Follicle quantification

Follicle quantification was performed on 5 μm ovarian sections stained with hematoxylin or subjected to immunostaining as indicated. To avoid repeated counting of the same follicle, every fifth serial section throughout the ovary was analyzed. Images were acquired using identical microscope settings for all experimental groups. Follicles were classified according to established morphological criteria. Primordial follicles were identified by the presence of an oocyte surrounded by a single layer of flattened pre-granulosa cells. Primary follicles were defined as oocytes enclosed by a single layer of cuboidal granulosa cells, whereas secondary follicles contained two or more layers of granulosa cells surrounding the oocyte. In addition, TAp63α was immunostained specifically to detect the primordial, primary and secondary follicles. Only follicles containing a clearly visible oocyte nucleus were included in the analysis to prevent duplicate counting. For each ovary, follicle numbers from all analyzed sections were summed and used to generate a single biological replicate. Follicle counting was performed in a blinded manner using Zeiss software Zen3.7. Data are presented as mean ± standard deviation (SD) from at least three independent biological replicates.

### Image acquisition and quantification

Fluorescence images were acquired using identical microscope settings for all experimental groups. Quantitative image analysis was performed using ZEN software (Carl Zeiss,blue edition 3.0, Germany). Bright-field images for all experimental groups were captured using the Nikon Eclipse TS2 microscope under identical imaging conditions. Exposure settings and acquisition parameters were kept constant across all experimental groups to ensure uniformity. IHC satined images for γH2AX, TAp63α, phospho-TAp63α, WEE1, phospho-WEE1, KU80, and XPA staining were captured using a Nikon fluorescence microscope under identical acquisition settings for all groups.

For fluorescence intensity measurements, regions of interest (ROIs) were manually drawn around individual oocytes or follicles, and mean fluorescence intensity values were determined following background subtraction. The same acquisition and analysis parameters were applied to all samples within an experiment. For TAp63α, MVH, phospho-CDK1(T14,Y15), phospho-CDK1(T161),RAD51, phospho-RAD51, and Tunnel, nuclear foci were manually quantified in individual oocytes. Only follicles containing a clearly visible oocyte nucleus were included in the analysis. Multiple follicles from at least three independent ovarian sections were analyzed for each biological replicate. A minimum of 30–50 oocytes per treatment group were evaluated unless otherwise indicated.

For marker-positive follicle analysis, the percentage of positive follicles was calculated relative to the total number of follicles examined per section. Quantification of fluorescence intensity, foci number, and marker-positive follicles was performed in a blinded manner, with investigators unaware of treatment allocation during image acquisition and analysis.

### Western blot

Goat ovaries were homogenized in lysis buffer (25 mM Tris-HCl, pH 7.5; 150 mM NaCl; 1% Triton X-100; 1% SDS; 100 mM PMSF; and 1× protease inhibitor cocktail) following the protocol described by Kumar et al. (2024). The lysates were centrifuged at 12,000 rpm for 10 minutes at 4°C, and the supernatant was collected for protein analysis. Protein samples were resolved by SDS-PAGE and transferred onto PVDF membranes. Membranes were blocked with 5% skimmed milk in TBST (Tris-buffered saline with 0.1% Tween-20) for 1 hour at room temperature and then incubated overnight at 4°C with the following primary antibodies: mouse anti-phospho-γH2A.X (05-63-I; 1:1000), mouse anti-PCRPChk2-1A9 (DSHB-S1-1758; 1:1000), mouse anti-phospho-Chk1/2 (9931T; 1:1000), rabbit anti-MVH (ab13840; 1:1000), rabbit anti-CDK1 (PAB888Mu01; 1:500), mouse anti-TAp63α (CM163B, 1:1000), rabbit anti-TAp63α (S160/162) (4981S; 1:1000), rabbit anti-TAp63α (S 455) (STJ90859; 1:1000),rabbit anti- phospho-CDK1 (T14, Y15) (44-686G; 1:500), rabbit anti- phospho-CDK1 (T161) (PA5-117191; 1:500), rabbit anti CK1α (DF-3185; 1:1000),rabbit anti-KU80(2180; 1:500), rabbit anti-XPA (PA5-119824; 1:500), anti-RAD51 (PC130; 1:1000), rabbit anti-phospho-Rad51(T309) (BS-5677R; 1:500) and mouse anti-β-Actin (SC-47778; 1:1000). Following primary antibody incubation, membranes were washed three times with TBST (5 minutes each), then incubated with HRP-conjugated secondary antibodies—goat anti-mouse IgG (31430, 1:10,000) and goat anti-rabbit IgG (31460, 1:5,000), for 2 hour at room temperature. After secondary incubation, membranes were washed three times with TBST, and protein bands were visualized using enhanced chemiluminescence detection.

### Western blot quantification

Western blot band intensities were quantified using ImageJ software (National Institutes of Health, Bethesda, MD, USA). For each blot, the integrated density of γH2AX, TAp63α , phospho-TAp63α, WEE1, phospho-WEE1, CDK1, phospho-CDK1 (Thr14/Tyr15 and Thr161), RAD51, phospho-RAD51, Ku80, XPA and other indicated proteins was measured following background subtraction. Protein expression levels were normalized to β-actin, which served as the loading control. Relative protein abundance was calculated by comparing normalized values to the corresponding vehicle control group. Quantification was performed using at least two independent biological replicates, and the resulting normalized values were used for statistical analyses. Data are presented as mean ± standard deviation (SD).

### TUNEL Assay

TUNEL staining was performed using the In-Situ Cell Death Detection Kit, Fluorescein (Cat. No. 11684795910). Mice ovary sections were fixed and processed for TUNEL staining. Antigen retrieval was performed using antigen retrieval buffer for 10 minutes to improve accessibility of fragmented DNA ends. Following retrieval, samples were washed with TBST. To minimize non-specific binding, samples were blocked using 10% ADB blocking solution for 15 minutes, repeated twice at room temperature. The TUNEL reaction mixture was prepared freshly according to the manufacturer’s protocol by mixing the enzyme solution and label solution in the recommended ratio. The reaction mixture was then added to completely cover the samples, followed by incubation in a humidified chamber at 37°C for 60 minutes in the dark. After incubation, samples were washed with PBS to remove excess reagent. Nuclei were counterstained using DAPI for visualization of nuclei. Finally, stained samples were mounted using DPX and observed under a fluorescence microscope. TUNEL-positive follicles showing green fluorescence were considered apoptotic. Quantification was performed by calculating the percentage of TUNEL-positive follicles relative to total follicles per section.

### Mouse experimental design

#### CDK1 inhibition in neonatal mice

To determine whether inhibition of CDK1 affects the establishment and maintenance of the ovarian reserve, neonatal female C57BL/6 mice were administered the CDK1 inhibitor RO-3306 by intraperitoneal injection. Mice received vehicle control or RO-3306 at the indicated doses (10 or 20 mg/kg body weight) once daily for 5 consecutive days starting at postnatal day 7 (P7). Ovaries were collected 24 h after the final injection, fixed in 10% formalin, processed for histological analysis, and subjected to immunofluorescence staining and follicle quantification.

#### Cisplatin-induced DNA damage model

To induce DNA damage in primordial follicle oocytes, neonatal female C57BL/6 mice were administered cisplatin by intraperitoneal injection at doses of 1, 2.5, or 5 mg/kg body weight. Control animals received vehicle alone. Ovaries were harvested 48 h after treatment and processed for immunofluorescence, immunohistochemistry, western blotting, and follicle analysis to assess DNA damage and follicular survival.

#### Effect of CDK1 inhibition on DNA damage repair

To evaluate the role of CDK1 during oocyte DNA repair, neonatal female mice were treated with cisplatin alone or in combination with the CDK1 inhibitor RO-3306. RO-3306 was administered before cisplatin treatment at the indicated dose. Ovaries were collected 24 h after treatment and analyzed for γH2AX, RAD51, phospho-RAD51, Ku80, and XPA expression by immunostaining and western blotting.

#### Role of Chk2 in DNA damage-induced CDK1 activation

To investigate the involvement of Chk2 signalling in DNA damage-induced CDK1 activation, neonatal female mice received cisplatin in the presence or absence of the Chk2 inhibitor BML-277. BML-277 was administered before cisplatin treatment at the concentrations indicated in the figure legends. Ovaries were collected 24 h after treatment and analyzed for Chk2 activation, WEE1 expression, CDK1 phosphorylation status, and DNA damage response markers.

#### Assessment of meiotic resumption following CDK1 activation

To determine whether DNA damage-induced CDK1 activation promotes meiotic resumption, fully grown germinal vesicle (GV)-stage oocytes were isolated from ovaries of treated and control mice and cultured in vitro under maturation conditions. After culture, oocytes were fixed and stained for phospho-H3 (Ser10), α-tubulin, and DNA. Meiotic progression was assessed by evaluating germinal vesicle breakdown (GVBD), spindle formation, chromosome condensation, and metaphase I entry.

#### Mice oocyte collection and In vitro maturation

Female mice were randomly assigned to three experimental groups: Vehicle Control (VC), Cisplatin (Cis), and Cisplatin plus CDK1 inhibitor (Cis + CDK1i) (n = 2 animals per group). Treatments were administered by intraperitoneal injection. Twenty-four hours after treatment, mice were euthanized, and ovaries were collected in M2 medium. Fully grown oocytes were isolated by mechanically puncturing ovaries with sterile needles in M2 medium under a stereomicroscope. Isolated oocytes were subsequently divided into two experimental groups. One group was immediately fixed in 4% paraformaldehyde (PFA) for assessment of meiotic status at the time of collection. The second group was cultured in in vitro maturation (IVM) medium under standard culture conditions for 8 h and subsequently fixed in 4% PFA for further analysis.

#### Oocyte immunofluorescence staining

Fixed oocytes were washed in TBST (Tris-buffered saline containing Tween-20) and blocked in antibody dilution buffer (ADB) for 15 min at room temperature. Following blocking, oocytes were incubated overnight at 4°C with mouse anti-α-tubulin (F2168, Sigma-Aldrich; 1:200) and rabbit anti-phospho-histone H3 (Ser10) (06-570, Millipore; 1:100) antibodies diluted in ADB. Antibody incubations were performed in 35-mm culture dishes in a final volume of 100 μL. The following day, oocytes were washed three times with TBST for 5 min each and re-blocked in ADB for 15 min at room temperature. Oocytes were then incubated with Alexa Fluor 555-conjugated goat anti-rabbit secondary antibody (A32732, Thermo Fisher Scientific; 1:2000) for 1 h at 37°C in the dark. After secondary antibody incubation, oocytes were washed three times with TBST for 5 min each. Stained oocytes were mounted on glass slides using ProLong Diamond Antifade Mount (P36970, Thermo Fisher Scientific) containing DAPI (0.1 μg/mL) and covered with glass coverslips. Fluorescence images were acquired using a Carl Zeiss, Axio Observer 7 Motorized fluorescence microscope and analyzed for spindle organization, chromosome condensation, and meiotic progression.

#### Statistical Analysis

All quantitative data are presented as mean ± standard deviation (SD) from at least three independent biological replicates unless otherwise stated. Statistical analyses were performed using GraphPad Prism (version 10.0; GraphPad Software). Comparisons between two groups were analyzed using an unpaired two-tailed Student’s *t*-test. Differences were considered statistically significant at *P* < 0.05. For western blot analyses, band intensities were quantified using ImageJ software (National Institutes of Health, Bethesda, MD, USA) and normalized to β-actin levels. The normalized values were used for statistical comparisons. Immunofluorescence, immunohistochemistry, TUNEL, and follicle quantification analyses were performed using images obtained from at least three independent biological replicates. The number of samples analyzed for each experiment is indicated in the corresponding figure legends.

## Methods contact

Further information and requests for resources and reagents should be directed to the lead contact, H.B.D. Prasada Rao

## ACKNOWLEDGMENTS

We thank the NIAB core Microscope and small animal facility. A.K.E was supported by UGC SRF. L.K.S. was supported by DBT SRF. A.K. was supported by DBT SRF, P.K. was supported by UGC JRF. This work was supported by the NIAB core grant C0031 and the DBT grant BT/PR46563/AAQ/1/868/2022 awarded to H.B.D.P.R.

## AUTHOR CONTRIBUTIONS

A.K.E, L.K.S and H.B.D.P.R. conceived the study and designed the experiments. A.M., A.K., P.K., P.K., L.K.S, A.S., S.S, S.Y., and H.B.D.P.R. performed the experiments and analyzed the data. H.B.D.P.R., L.K.S and A.K. wrote the manuscript with inputs and edits from all authors.

## Conflict of interest

The authors declare no competing interests. None of the material reported in this manuscript has been published or made available online, nor is it under consideration elsewhere.

---

**Model:**

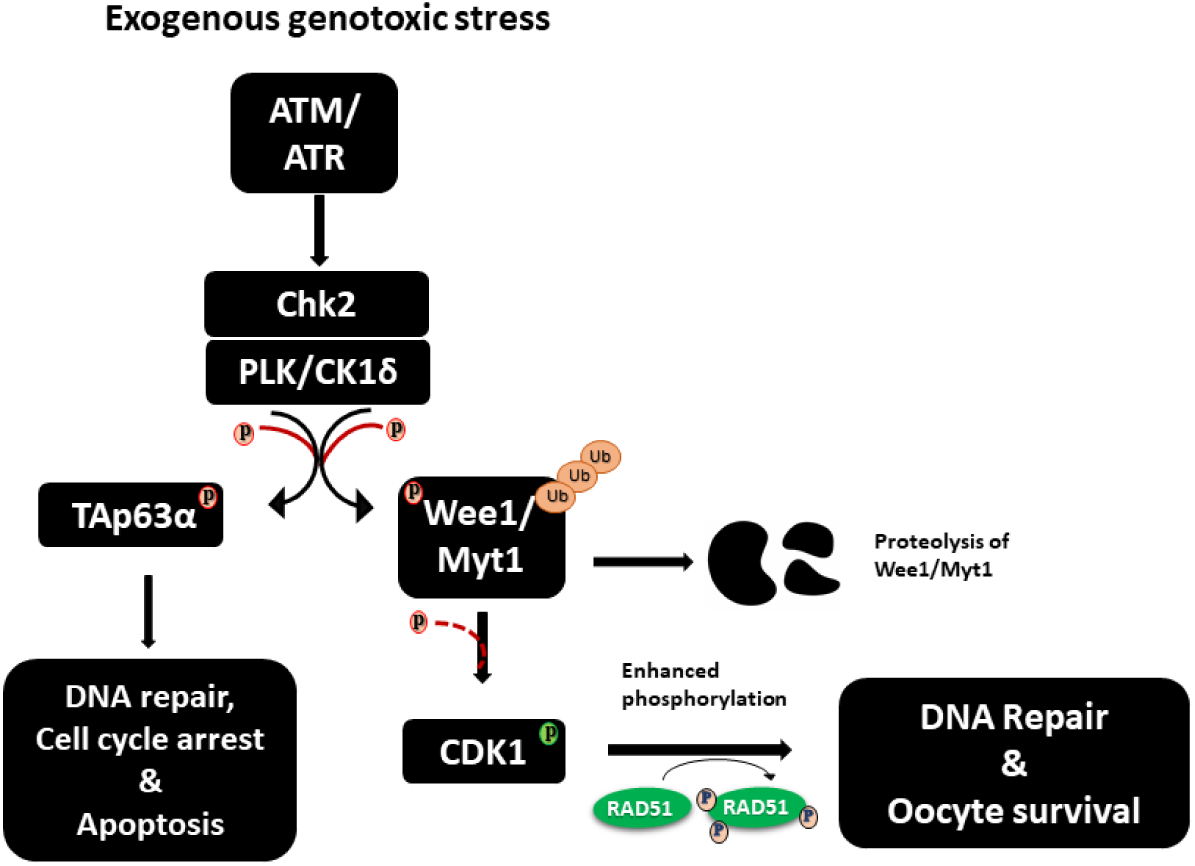

## Proposed model of oocyte response to exogenous DNA damage

Exposure of oocytes to exogenous DNA damage activates the DNA damage response (DDR), which is initiated by the ATM/ATR signaling pathway. Activation of the downstream checkpoint kinase Chk2 promotes phosphorylation of TAp63α, a key guardian of oocyte quality control. In response to severe or persistent DNA damage, activated TAp63α induces the expression of pro-apoptotic factors, leading to the elimination of damaged oocytes and preventing the transmission of genomic abnormalities.

In parallel, our findings suggest that Chk2 participates in the regulation of DNA repair through modulation of the WEE1–CDK1 checkpoint pathway. DNA damage-induced Chk2 activation is associated with reduced WEE1 activity and increased CDK1 activation, as evidenced by decreased inhibitory phosphorylation (T14/Y15) and increased activating phosphorylation (T161) of CDK1. Activated CDK1 promotes RAD51 phosphorylation and is associated with enhanced HR-mediated DNA repair. In contrast, inhibition of CDK1 suppresses RAD51 activation and is accompanied by increased expression of the NHEJ and NER repair markers KU80 and XPA. Together, these findings support a model in which Chk2 coordinates two complementary responses to DNA damage in oocytes: activation of the TAp63α-dependent quality control pathway and regulation of CDK1-dependent DNA repair mechanisms. This dual response may enable oocytes to either repair manageable DNA lesions or undergo elimination when genomic damage exceeds the capacity for repair, thereby preserving ovarian reserve quality and genomic integrity.

## Supplementary Figures

**Supplementary Figure 1:**
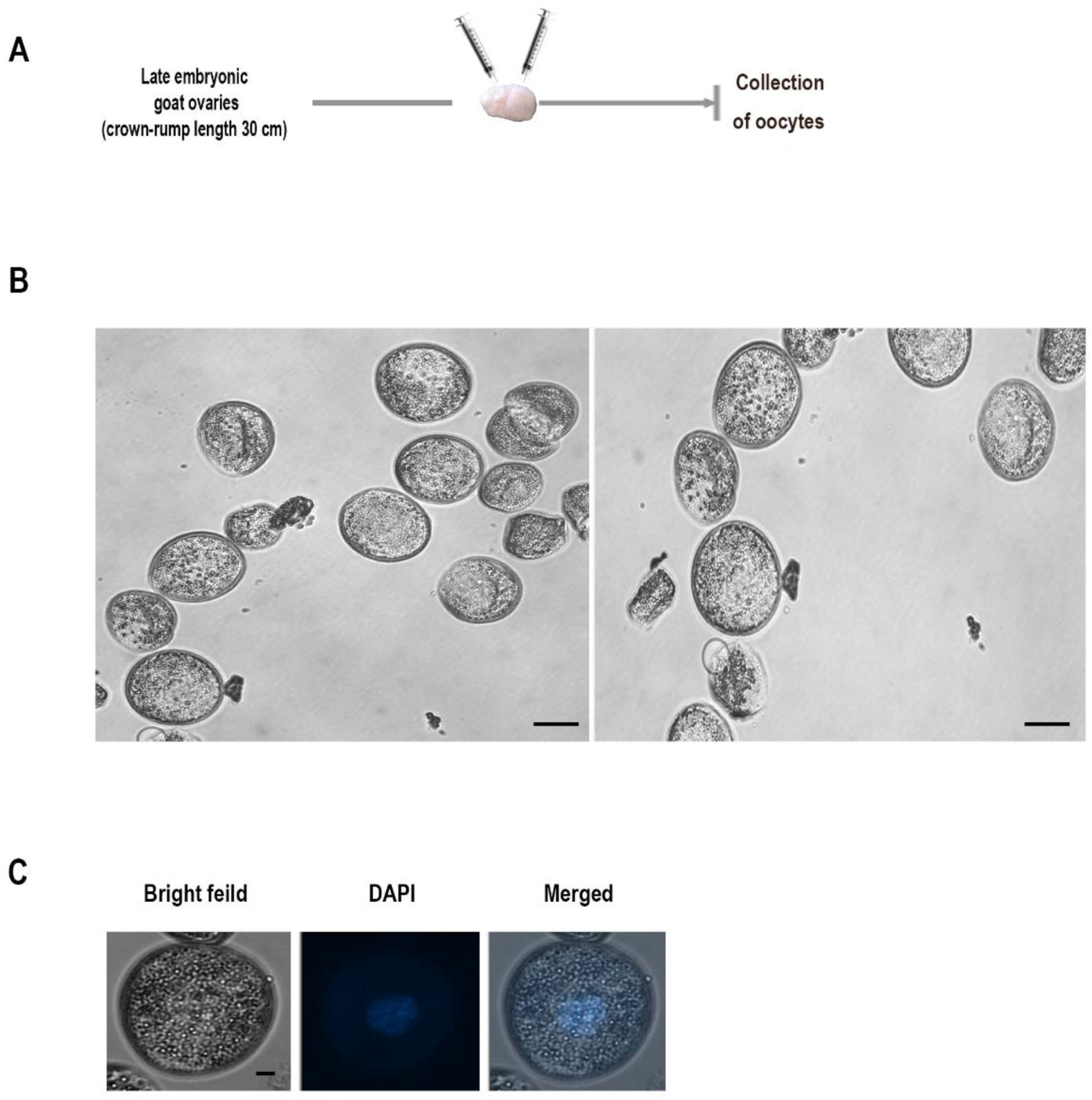
Isolation and characterization of dictyate-stage oocytes from goat fetal ovaries. (A) Schematic overview of the procedure used for isolation of dictyate-stage oocytes from goat fetal ovaries. (B) Representative bright-field image showing a group of isolated dictyate-stage oocytes. Scale bar = 50 μm. (C) Representative images of individual dictyate-stage oocytes shown in bright-field and DAPI channels. Merged images are also presented to visualize nuclear morphology. Scale bar = 10 μm.

**Supplementary Figure 2:**
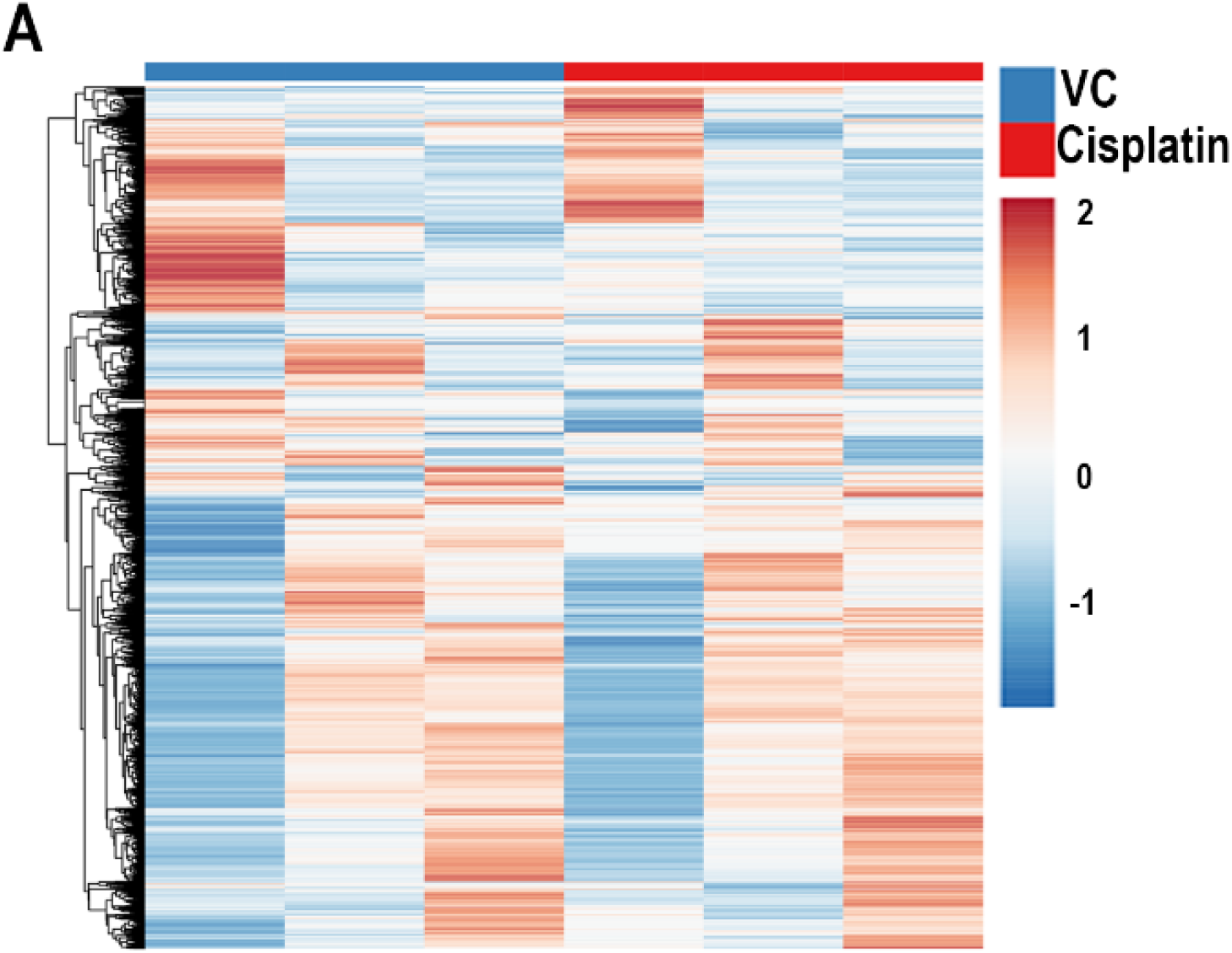
(A) Heatmap depicting the differential expression of proteins in vehicle control (VC) and Cisplatin-treated ovaries, highlighting up- and down-regulated candidates.

**Supplementary Figure 3:**
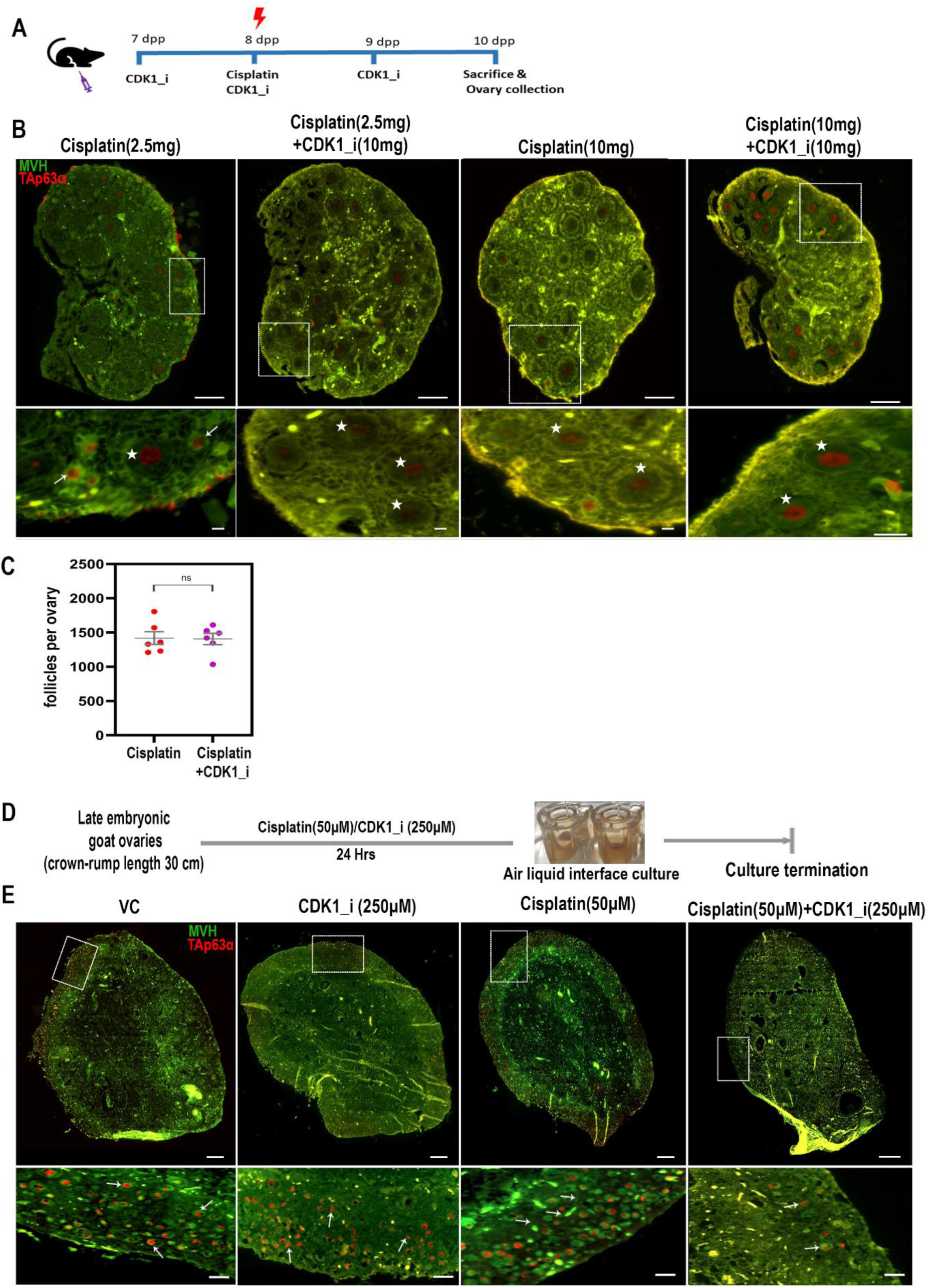
Effect of CDK1 inhibition on ovarian pool following cisplatin treatment. (A) Schematic overview of the in vivo experimental design. Neonatal mice were treated with cisplatin alone or cisplatin in combination with a CDK1 inhibitor by intraperitoneal injection. (B) Representative immunofluorescence images of ovarian sections from cisplatin-treated and cisplatin plus CDK1 inhibitor-treated mice stained for MVH (green) and TAp63α (red). White boxes indicate regions shown at higher magnification in the lower panels. Arrows indicate TAp63α-positive primordial follicles, and asterisks indicate TAp63α-positive secondary follicles. Scale bars: 100 μm (upper panels) and 10 μm (lower panels). (C) Quantification of total follicle numbers per ovary in cisplatin-treated and cisplatin plus CDK1 inhibitor-treated mice (n = 6 ovaries per group). (D) Schematic overview of the ex vivo experimental design using cultured late embryonic goat ovaries. Ovaries were treated with Vehicle Control (VC), CDK1 inhibitor alone (CDK1i), Cisplatin (Cis), or Cisplatin plus CDK1 inhibitor (Cis + CDK1i). (E) Representative immunofluorescence images of goat ovarian sections from the indicated treatment groups stained for MVH (green) and TAp63α (red). White boxes indicate regions shown at higher magnification in the lower panels. Arrows indicate TAp63α-positive follicles. Scale bars: 200 μm (upper panels) and 50 μm (lower panels). Data are presented as mean ± SD. Statistical significance was determined using an unpaired two-tailed Student’s *t*-test. ns, not significant.

**Supplementary Figure 4:**
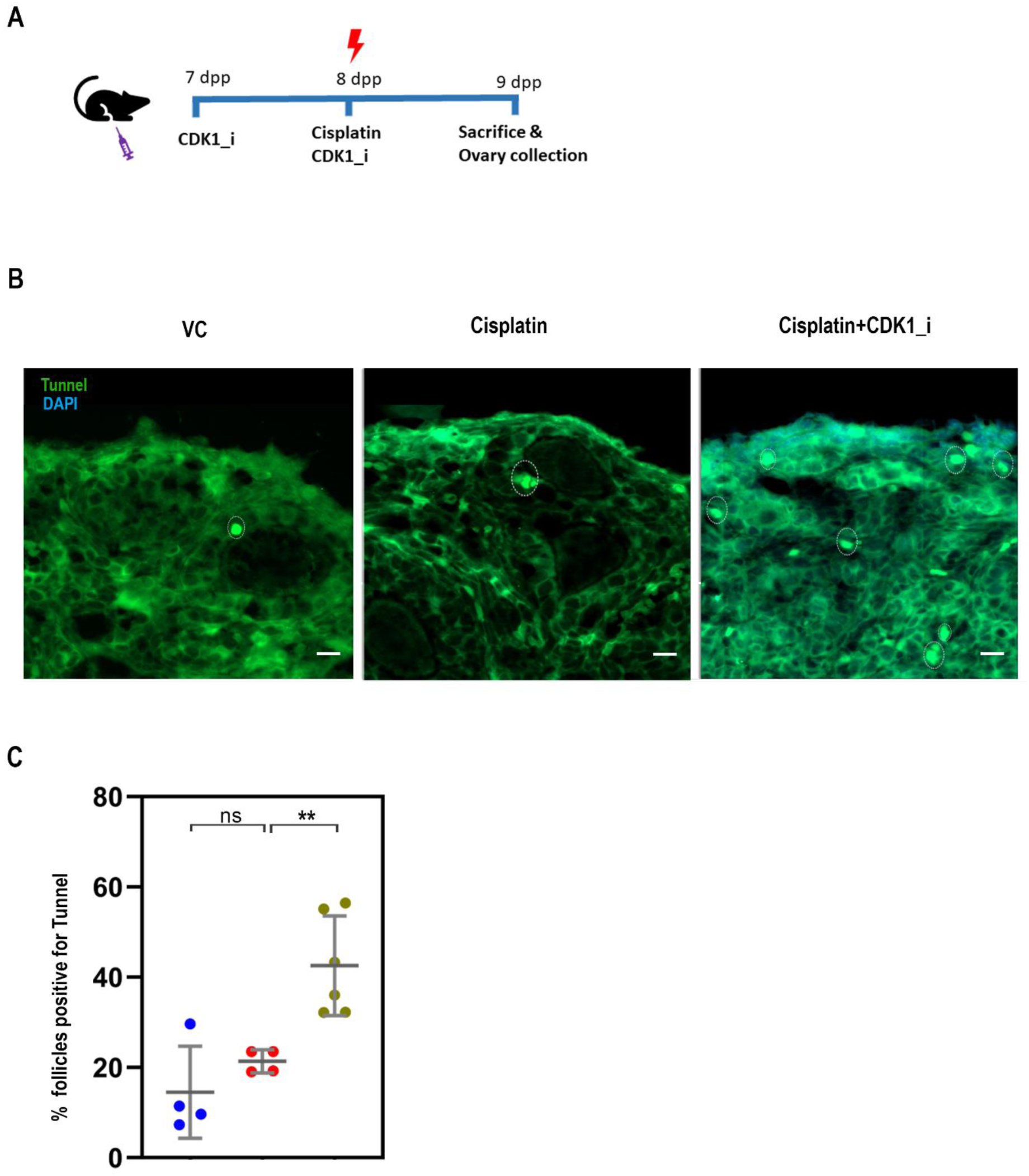
TUNEL analysis of apoptosis in ovarian follicles following cisplatin treatment and CDK1 inhibition. (A) Schematic overview of the experimental design. Neonatal mice were assigned to three treatment groups: Vehicle Control (VC), Cisplatin (Cis), and Cisplatin plus CDK1 inhibitor (Cis + CDK1i). Treatments were administered by intraperitoneal injection. (B) Representative TUNEL staining of ovarian sections from the indicated treatment groups. TUNEL-positive cells are shown in green, and nuclei are counterstained with DAPI (blue). Scale bar = 10 μm. (C) Quantification of TUNEL-positive follicles per ovarian section in the indicated treatment groups (n = 4 sections per group). Data are presented as mean ± SD. Statistical significance was determined using unpaired two-tailed Student’s *t*-tests for predefined pairwise comparisons between experimental groups. Significance levels are indicated as **P ≤ 0.01, and ns, not significant.

**Supplementary Figure 5:**
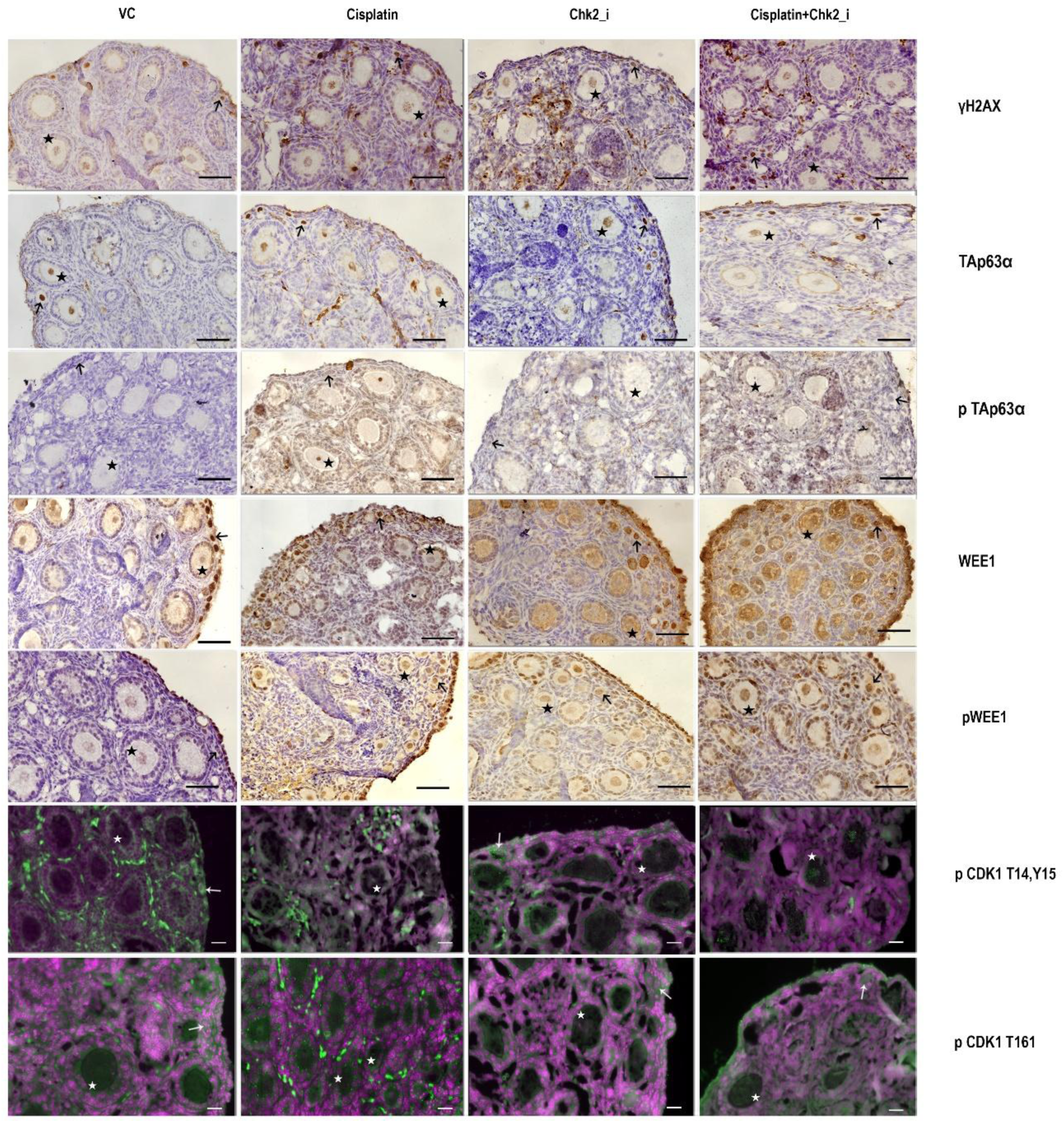
Enlarged views of representative follicles from Figure 6. Representative enlarged images of primordial and secondary follicles cropped from the ovarian sections shown in Figure 6 for γH2AX, TAp63α, phospho-TAp63α (p-TAp63α), WEE1, phospho-WEE1 (p-WEE1), phospho-CDK1 (T14/Y15), and phospho-CDK1 (T161) staining. Arrows indicate primordial follicles, and asterisks indicate secondary follicles. Scale bars: 50 μm for immunohistochemistry (IHC) images and 20 μm for immunofluorescence (IF) images.

**Supplementary Figure 6:**
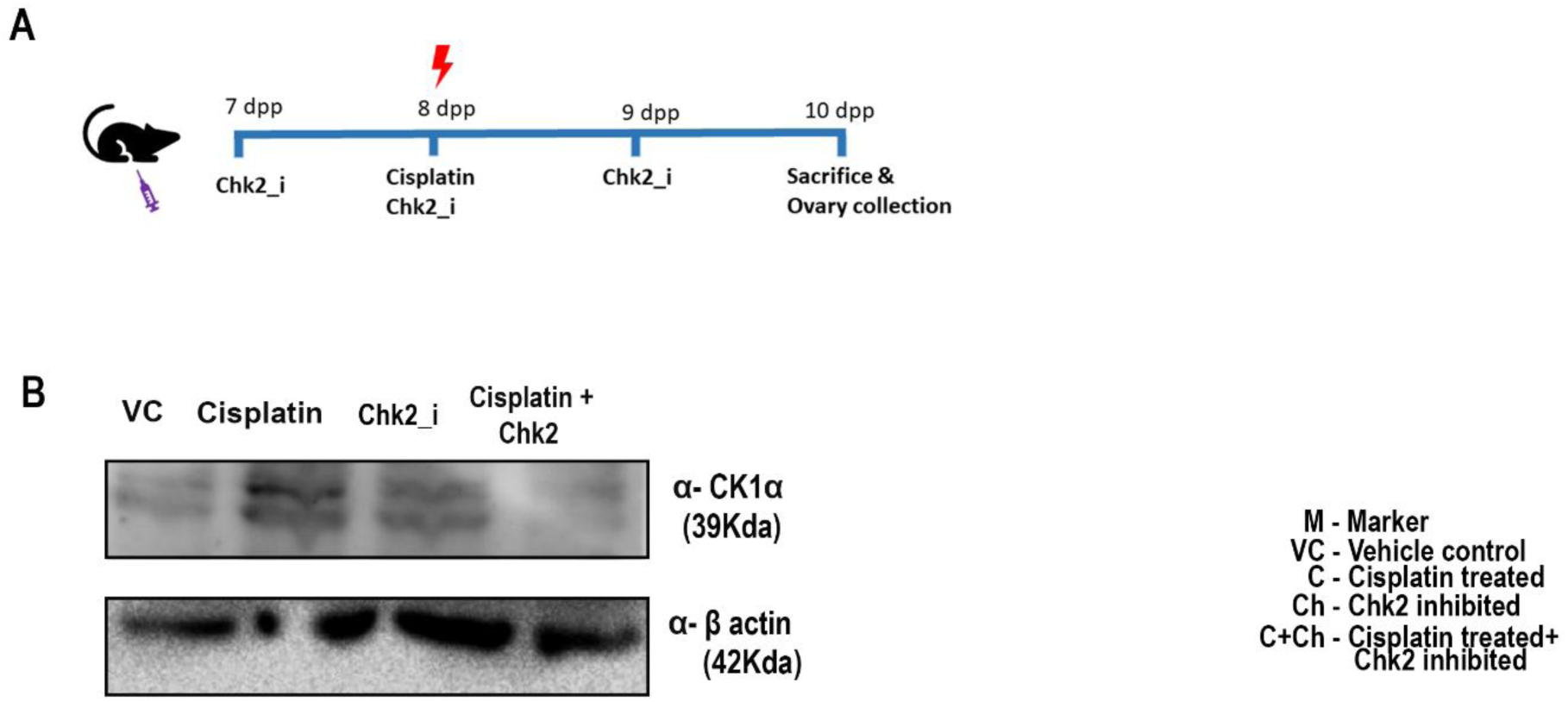
(A) Schematic overview of the experimental design. Neonatal mice were assigned to four treatment groups: Vehicle Control (VC), Chk2 inhibitor alone (Chk2_i), Cisplatin (Cis), and Cisplatin plus Chk2 inhibitor (Cis + Chk2_i). Treatments were administered by intraperitoneal injection. (B) Representative western blots of ovarian lysates from the indicated treatment groups showing expression of CK1α and β-actin as a loading control. M indicates marker, VC-vehicle control, C- Cisplatin-treated ovary, Ch- Chk2 inhibitor and C+Ch- cisplatin combined with Chk2 inhibitor.

**Supplementary Figure 7:**
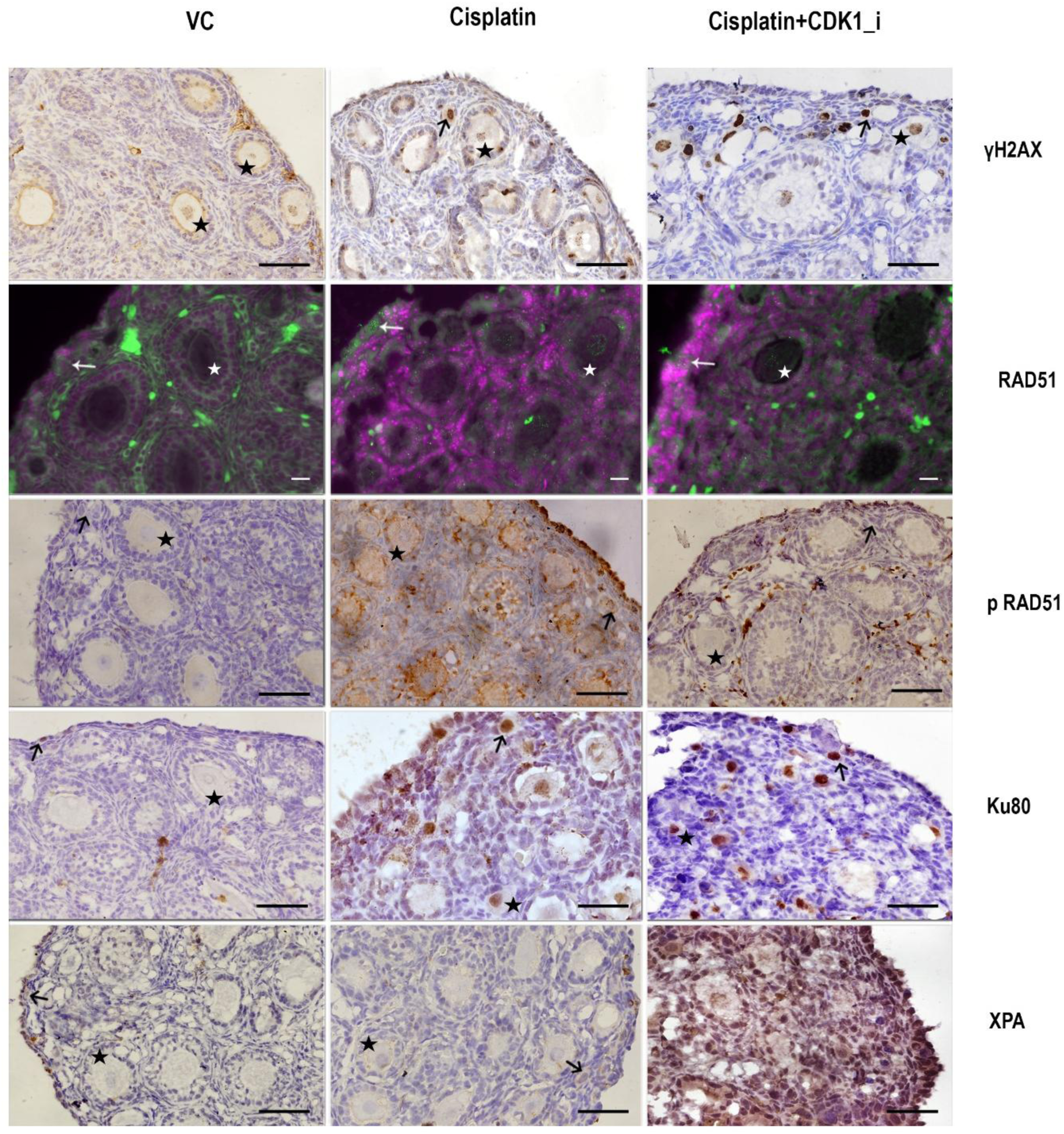
Enlarged views of representative follicles from Figure 7. Representative enlarged images of primordial and secondary follicles cropped from the ovarian sections shown in Figure 7 for γH2AX, RAD51, phospho-RAD51 (p-RAD51), KU80, and XPA staining. Arrows indicate primordial follicles, and asterisks indicate secondary follicles. Scale bars: 50 μm for immunohistochemistry (IHC) images and 20 μm for immunofluorescence (IF) images.

**Supplementary Figure 8:**
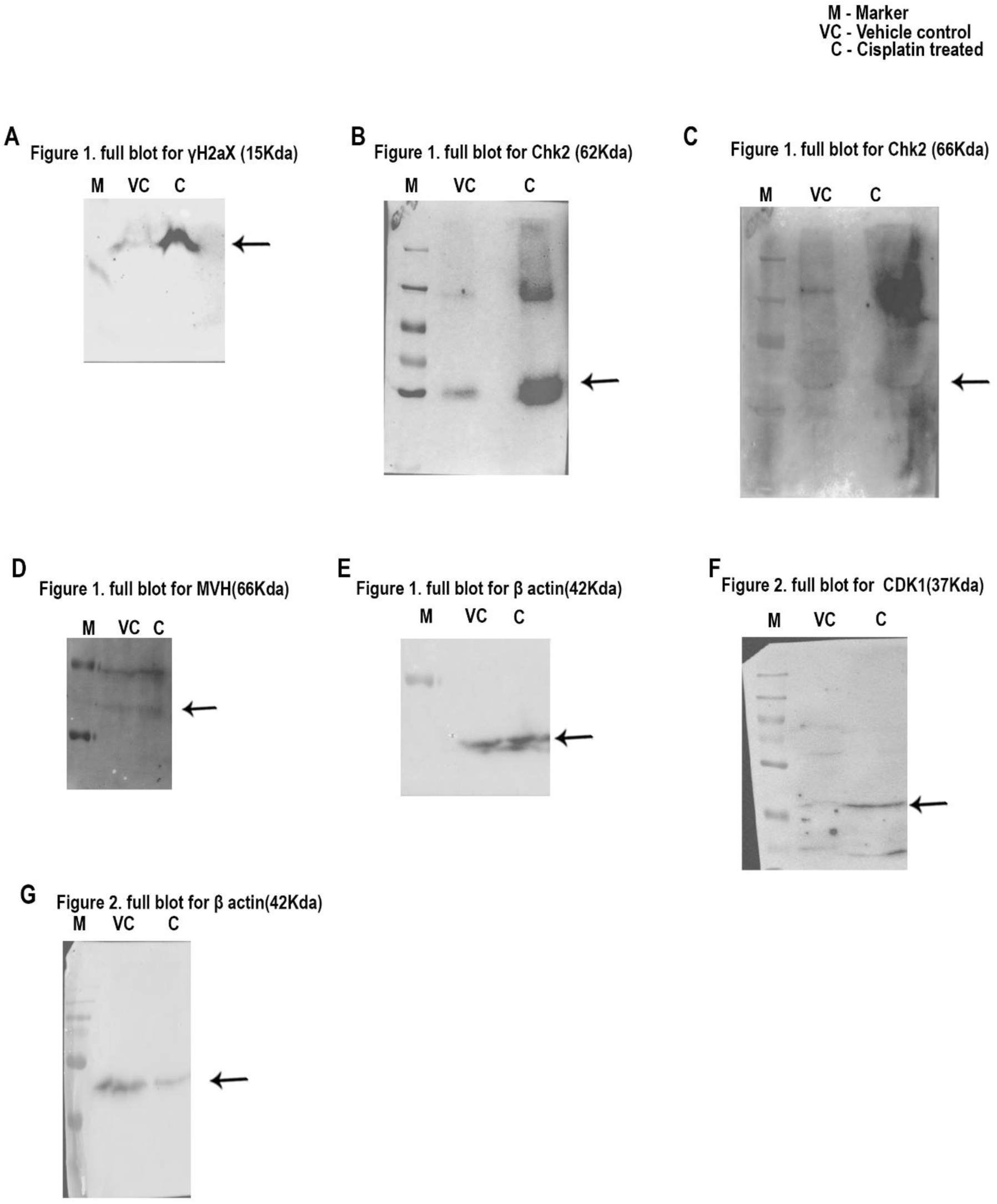
Uncropped full Western blots. (A-G) Full western blot images of γH2aX, Chk2, pChk2, MVH, β-Actin, CDK1 and β-Actin in VC and DNA-damaged ovaries. M indicates marker, VC- vehicle control and C- Cisplatin treated ovary.

**Supplementary Figure 9:**
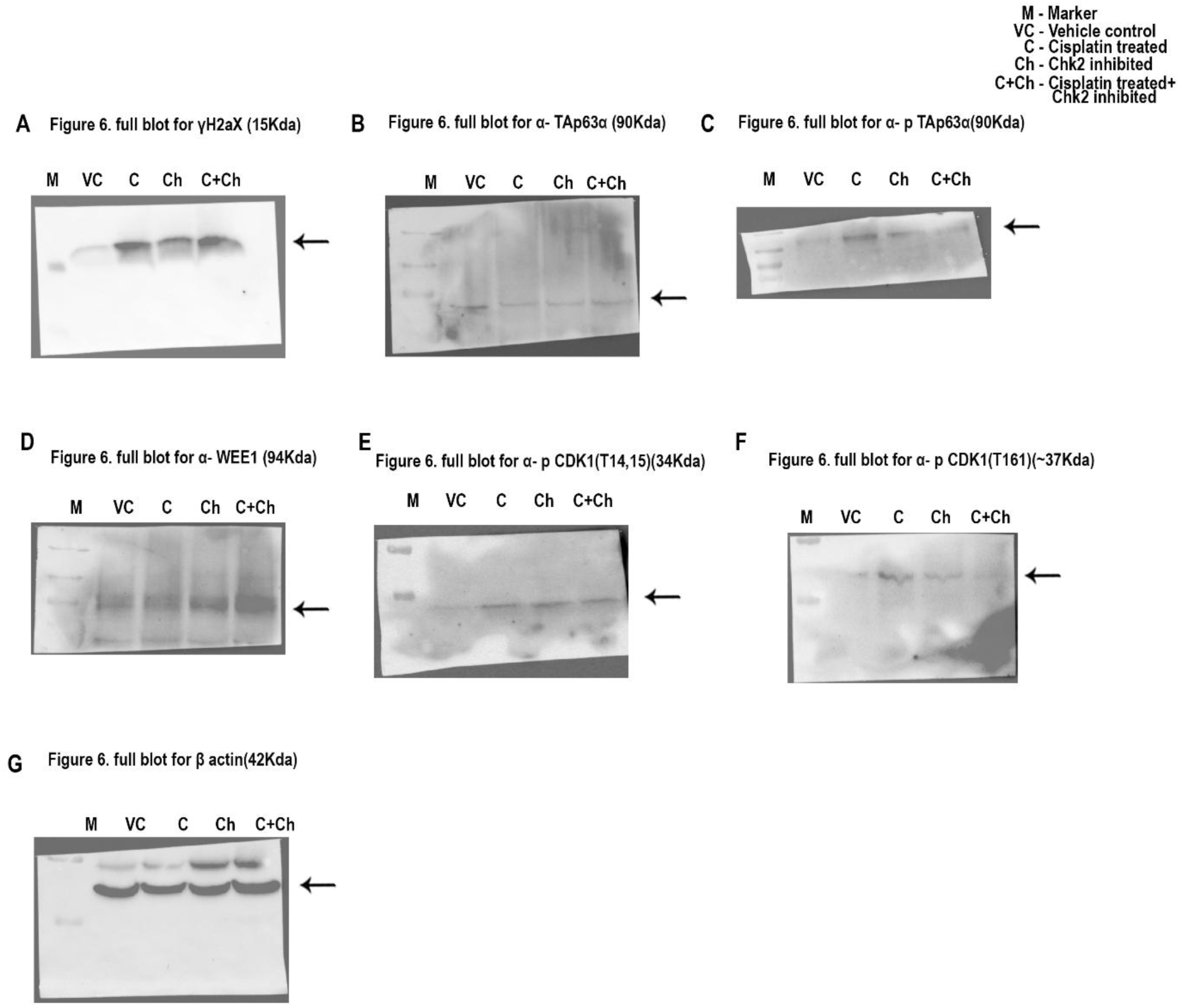
Uncropped full Western blots. (A-H) Full western blot images of γH2AX, TAp63α, phospho-TAp63α, WEE1, phospho-CDK1 (T14/Y15), phospho-CDK1 (T161), and β-actin for the Vehicle Control (VC), Chk2 inhibitor alone, Cisplatin alone, and Cisplatin combined with Chk2 inhibitor experimental groups. M indicates marker, VC- vehicle control and C- Cisplatin treated ovary. M indicates marker, VC- vehicle control, C- Cisplatin-treated ovary, Ch- Chk2 inhibitor and C+Ch-cisplatin combined with Chk2 inhibitor. Arrow indicates the band.

**Supplementary Figure 10:**
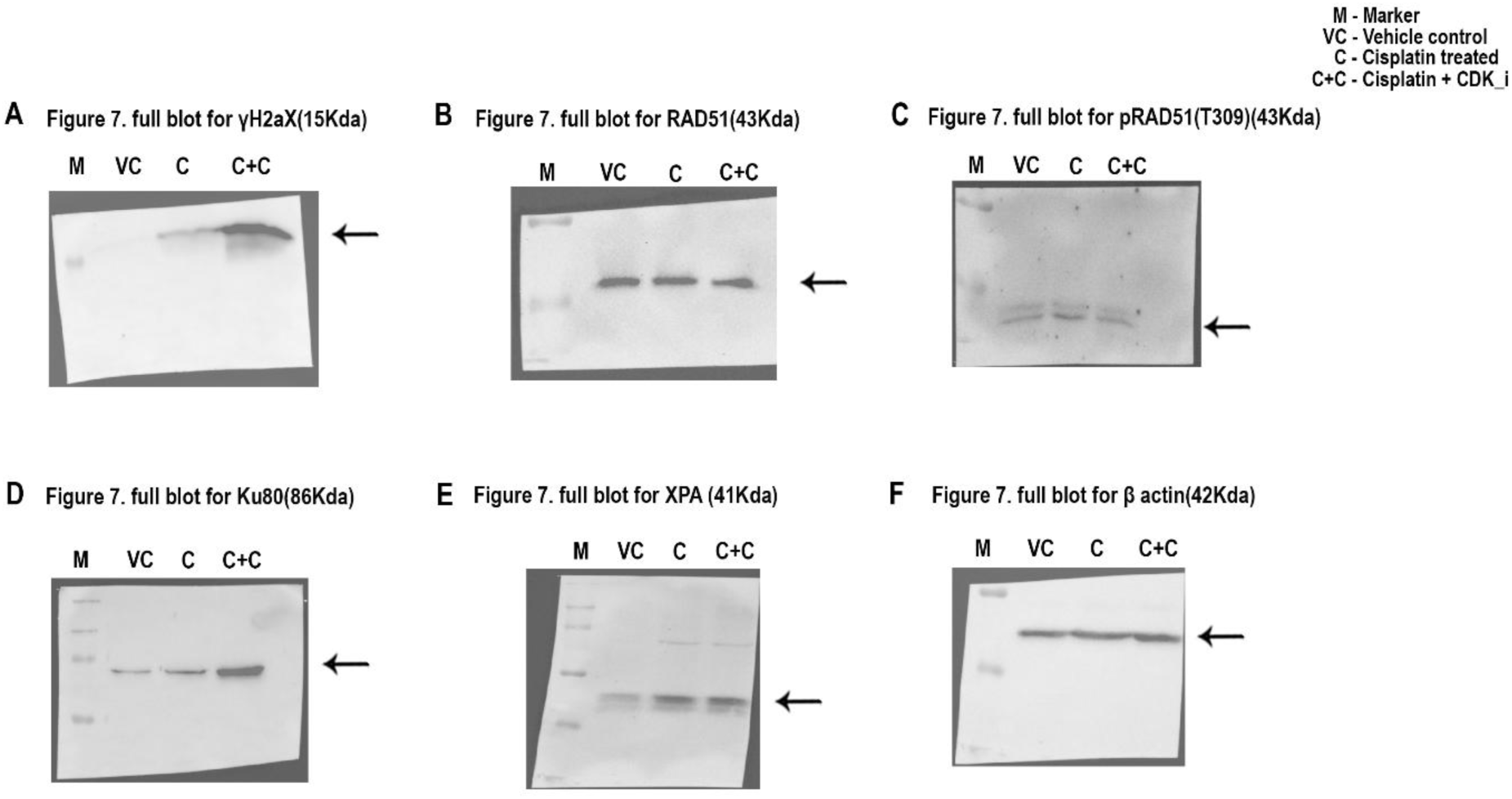
Uncropped full Western blots. (A-F) Full western blot images of γH2AX, RAD51, phospho-RAD51, KU80 and XPA and β-actin. M indicates marker, VC- vehicle control, C- Cisplatin-treated ovary, and C+C indicates Cisplatin treatment with CDK1 inhibitor.

## Notes

### Competing Interest Statement

The authors have declared no competing interest.

